# Influence of natural enemy specificity and functional response on victim coexistence

**DOI:** 10.64898/2025.12.04.692374

**Authors:** Dipanjana Dalui, Annette M Ostling, Colin T Kremer, Robert Bagchi

**Affiliations:** University of Nebraska, Lincoln; University of Texas, Austin; University of Connecticut, Storrs

**Keywords:** diversity maintenance, specialization, antagonistic interactions, host-pathogen

## Abstract

Natural enemies are thought to promote coexistence of competing victim species. Although existing theory suggests victim coexistence increases with enemy specialization, the dynamics and potential extinction of enemies is generally discounted. Where enemy dynamics have been considered, empirically atypical linear functional responses have been studied. These limitations could over-simplify inferences about enemy-mediated coexistence. We studied the dynamics of two competing victim species and two enemy species with a deterministic model. We derived equilibrium points, and used linear stability analysis, numerical simulations and Floquet theory to determine the influence of enemy specificity and non-linear functional responses on coexistence in this victim-enemy community. We found greater specificity could drive enemy equilibrium points to infeasible values. We found only accelerating enemy functional responses result in stable equilibrium point coexistence of otherwise equivalent competitor victims, in which case greater specificity results in greater stability. Linear and saturating responses produce complex dynamics (neutral or limit cycles, chaos) or extinction, with limit cycle stability highest at intermediate specificity. Our results indicate strict specificity may not maximize coexistence, and enemy functional response critically influences whether enemies promote victim coexistence. They highlight the need to incorporate enemy dynamics into the growing body of theory regarding enemy-mediated diversity maintenance.

## Introduction

A prominent explanation for the coexistence of many species with similar resource requirements is that natural enemies (including predators, herbivores and pathogens, hereafter enemies) regulate their victims’ populations and prevent competitive exclusion (Janzen 1970; Connell 1971; Levins 1979;Adler and Muller-Landau 2005; Bever et al. 2015). One mechanism for this enemy-mediated coexistence is that specialized enemies are attracted by dense aggregations of their victim species, reducing victim survival (creating “negative conspecific density dependence”). The mortality of densely aggregated species promotes recruitment of other victim species that might otherwise be out-competed (e.g. Janzen 1970; Connell 1971; Bever 1994; Grover and Holt 1998; Chesson 2018). An extensive body of research supports this hypothesis, including theoretical studies and simulations (Armstrong 1989; Adler and Muller-Landau 2005; Sedio and Ostling 2013; Levi et al. 2019; Chisholm and Fung 2020), and empirical studies spanning both tropical and temperate ecosystems (Harms et al. 2000; Packer and Clay 2000; Lambers et al. 2002; Johnson et al. 2012 Bagchi et al. 2014, 2010; Comita et al. 2014). A large proportion of this work has concentrated on plant communities, with a focus on testing the importance of enemy-mediated coexistence. This work has indicated that fungal and oomycete pathogens are important drivers of conspecific-density dependence in tree seedling assemblages (Augspurger 1984; Augspurger and Wilkinson 2007; Bagchi et al. 2014, 2010; Mangan et al. 2010), with evidence that insect herbivores may also contribute (Sullivan 2003; Forrister et al. 2019). However, a number of theoretical and empirical questions remain to be answered to fully delineate the role of enemies in the maintenance of diversity of their hosts plants. In this study, we focus on two characteristics of the enemy-victim interaction that we identified to be crucial for coexistence; (1) the specificity of the enemy-victim interaction, and (2) the functional response of the enemies to victim densities. Specificity of enemies is crucial for generating conspecific density dependence; without it, enemies would not respond to high densities of a single victim species (at least not without environmental variation and partitioning simultaneously at play; Mordecai 2015). An overwhelming proportion of theoretical studies to date have assumed, explicitly or implicitly, that enemies specialize on a single victim species (Adler and Muller-Landau 2005; Levi et al. 2019). However, complete host-specificity appears rare in nature, with many fungi and insects able to attack many, usually related, species even in the tropics (Ødegaard et al. 2000; Novotny and Basset 2005; Ødegaard et al. 2005; Gilbert and Webb 2007; Novotny et al. 2010; Hersh et al. 2012; Spear and Broders 2021). This rarity of strict host-specific enemies in nature emphasizes the need for investigations of the relationship between the effectiveness of enemies at regulating their victims’ populations and their specificity. Kirchner and Roy (2002), also summarized in Bruijning et al. (2023), analyzed a two-host-two-pathogen model and suggested that host specificity facilitated coexistence (allowing invasion of a rare host into an equilibrium community controlled by a preferred-host/pathogen pair). However, coexistence of all four species in this model is not robust: the extinction of any single participant results in the loss of its associate (Kirchner and Roy 2002), highlighting the difficulties of using mutual invasibility to demonstrate coexistence in multi-trophic communities (Grainger et al. 2019). Using stochastic simulation models in which recruitment depended on the number of nearby adults sharing enemies (Adler and Muller-Landau 2005), Sedio and Ostling (2013) varied host-specificity on a continuous scale and indicated that, while the capacity for enemies to maintain victim diversity increases with host specificity, strict host specificity was not a prerequisite for this mechanism to substantially contribute to diversity. Using a deterministic model, Chesson (2018) also demonstrated that host-specificity increased the potential for victim species to coexist, quantified in terms of the contribution to the “stabilization” term of invasion growth rates. In essence, increasing host specificity of enemies decreases the niche overlap among enemies, where an array of enemies make up the niche axes. As a result, increasing population density of any victim species concentrates competition among conspecifics rather than heterospecifics and promotes coexistence. Because each enemy represents a new limiting factor, increasing numbers of host-specific enemies could, in theory, support the coexistence of an infinite number of victim species (Levin 1970).

The possibility of an unlimited number of coexisting victim species highlights an important issue with using specialized enemies to explain coexistence. Specialization on different victim species could support enemy species coexistence, potentially allowing an unlimited number of victim and enemy species to mutually support each other. However, although host-specialization provides several advantages to enemies (Futuyma and Moreno 1988), low densities of their victims could expose enemies to a high extinction risk (May 1991; Woolhouse 2001). Enemy dynamics, therefore, provide the limit to such infinite coexistence, as successive partitioning of resources eventually drives perilously low densities and high extinction risks. Enemy dynamics, however, have been largely overlooked during the development of theory of enemy-mediated coexistence of their victim species. Instead, the majority of models linking victim species coexistence to enemies have included enemy influence only implicitly, through a reduction in victim recruitment probability that depends on the number of individuals nearby and their relative similarity in enemies (Adler and Muller-Landau 2005; Sedio and Ostling 2013; Levi et al. 2019; Chisholm and Fung 2020). Although Chesson (2018) explicitly included enemy dynamics in a two-trophic model of species coexistence, minimum enemy population densities were bounded above zero in the absence of suitable victims, implicitly representing either the presence of additional victim species not explicitly modeled, or immigration or persistent life stages. Although these models suggest that the specificity of enemies increases the potential for victim-species coexistence, that conclusion may rely on this assumption that the enemy population has a source of population growth that does not require the presence of the focal victims in the model, which will allow it to persist in the absence of the victims and may also stabilize its population dynamics. Removing this assumption could reveal that overly specialized enemies may not have feasible equilibrium populations, and may have complex dynamics that could bring them near, or to, extinction. If an enemy does not have a feasible equilibrium, or should go extinct due to complex dynamics, its victim would be released from top-down regulation while other victims continue to be regulated by their own enemies or resources, potentially leading to mono-dominance in a community. Therefore, assessments of the role of host specialization in enemy-mediated species coexistence could benefit from explicit consideration of enemy dynamics.

Including enemy dynamics in models of victim coexistence requires assumptions about how enemy attack rate responds to victim density (i.e., the functional response, Holling 1965). For simplicity, most current models of enemy-victim interactions use linear relationships (i.e. “Type 1” functional responses) between victim densities and enemy attack rates (Hopkins et al. 2020; McCallum et al. 2001). However, existing empirical data indicate that attack by both pathogens and insects often responds non-linearly to increasing host density by saturating or accelerating as victim density increases (Bagchi et al. 2011; Orlofske et al. 2018; Hopkins et al. 2020). Embracing non-linear functional responses is necessary to adequately assess the potential for enemies to stabilize the coexistence of their victims, as linear functional responses, although simpler to analyze, lead to neutral cycles that are special to the case of exact linearity (Lotka 1925; Volterra 1927). The functional response of enemies can have important implications for enemy-victim dynamics (McCallum et al. 2001), and non-linear functional responses can arise from a variety of important details of natural dynamics. For example, adding exponents to the interaction equations can capture a myriad of instances like: enemy satiation, victim refuge, host switching, the spatial context of many enemy-victim interactions (Pascual et al. 2011), variation in interactions, overcompensating density dependence (Münkemüller et al. 2009; Bagchi et al. 2010), and cumulative enemy load. Incorporating diverse functional responses involving multiple species requires consideration of the complexity of functional responses of enemies who can consume multiple victim species, who would be expected to respond to the combined density of their victim species (Dobson 2004).

In this study, we explore the role of natural enemy specificity in allowing coexistence of their victims, while explicitly incorporating enemy population dynamics. We present a model of two enemy species and two victim species who are competing directly (as well as indirectly through the enemies). Our model flexibly characterizes enemy functional responses, allowing linear, saturating, and accelerating increases of attack rates in response to the combined density of victim species. We hypothesize that strict host specificity by enemies can lead to infeasible population sizes, and hence break-down of enemy-mediated coexistence. We test this by deriving expressions for equilibrium population sizes and determining the conditions in which they are feasible. We also analyze the stability of these four-species communities, determining the effect of functional response and enemy specificity together on stability. Doing so turns out to require not only analysis of linear stability around equilibrium points, but also the consideration of complex dynamics such as limit cycles and chaos, and the use of Flouquet theory to analyze the stability of limit cycles. To our knowledge, this study provides the most complete characterization to date of the impact two enemies can have on the coexistence of otherwise equivalent competing victims–a characterization that in particular considers the impact of both the degree of enemy specificity and the form of the functional response.

## Methods

Our model was built with the objective of investigating the interaction between victims (including prey and hosts) and their enemies (including predators, pathogens and herbivores) to answer two major questions: (1) how does specificity of enemies influence species coexistence of their competing victims? and (2) how is this coexistence influenced by the functional response of the enemies? We based our model on the classic Lotka-Volterra predator-prey model (Lotka 1925; Volterra 1927) with the following additions: (a) we added a second victim species and a second enemy species, (b) we added intraspecific and interspecific victim-victim competition (assumed equal in magnitude, to focus on victim coexistence generated by interactions with enemies), (c) we varied the preference of each natural enemy for each of the victims, and (d) we introduced a generalized functional response of enemies to the combined victim density.

In our model (1), two enemy species (*P*_*A*_; *P*_*B*_)consume two victim species (*H*_1_; *H*_2_), and the encounter leads to losses and gains in densities of the victim and enemy species respectively:

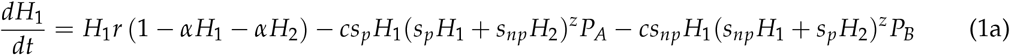

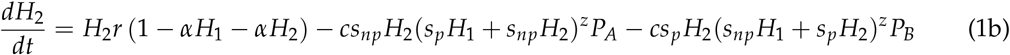

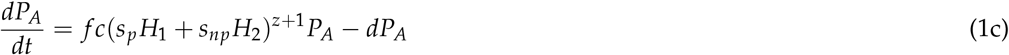

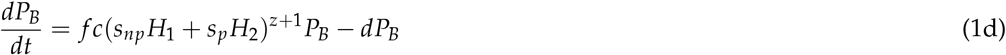

See Table 1 for model parameter definitions. The strength of each interaction is scaled by the susceptibility or vulnerability (henceforth, susceptibility) of the victims to each enemy species, where each enemy has a preferred victim (susceptibility =*s*_*p*_) and non-preferred victim (susceptibility =*s*_*np*_). We model the functional response of the enemies as a dependence on the density of its victims weighted by its preferences for them, allowing us to separately manipulate the functional response (through the form of this overall density-dependence) and the enemy specialization (through these weightings).

**Table 1:** Parameters and their definitions.

| Parameter | Definition |
| --- | --- |
| $r$ | intrinsic rate of population increase for victim species |
| $\alpha$ | competition coefficient for victim-victim interactions |
| $s_p$ | susceptibility (or vulnerability) of victim species to specialized enemy |
| $s_{np}$ | susceptibility (or vulnerability) of victim species to non-specialized enemy |
| $c$ | rate of contact (or attack) between a victim and its enemy |
| $f$ | rate of conversion of contacted victim into new enemy (also called fecundity) |
| $d$ | intrinsic death rate of enemy |
| $z$ | functional response of enemy to susceptibility-weighted summed victim densities |

The exponent *z* defines the functional response of enemies to the combined, susceptibility-weighted density of the two victim species (Fig. 1). *z* =0 represents the linear case in which enemy load increases linearly with victim density. When *z* > 0, enemy attack rate is an accelerating function of the susceptibility-weighted summed density of both victim species, i.e. the rate of attack of an enemy of its victim increases faster than linearly with victim density. This could occur, for example, because more densely packed victims are closer in space and may be substantially easier to disperse between. For −1 < *z* < 0 the relationship between attack rate and victim density is saturating, i.e. it levels off at high victim density. This could occur, for example, because the enemy is limited by some “handling time” in its interaction with victims, which could reflect the time to subdue and consume prey or to achieve a within-host population substantial enough to achieve transmission or dispersal to additional hosts. We do not consider the case when *z* < −1 because that would imply that enemies killed fewer victims as victim density increased, which is biologically implausible.

**Figure 1.**
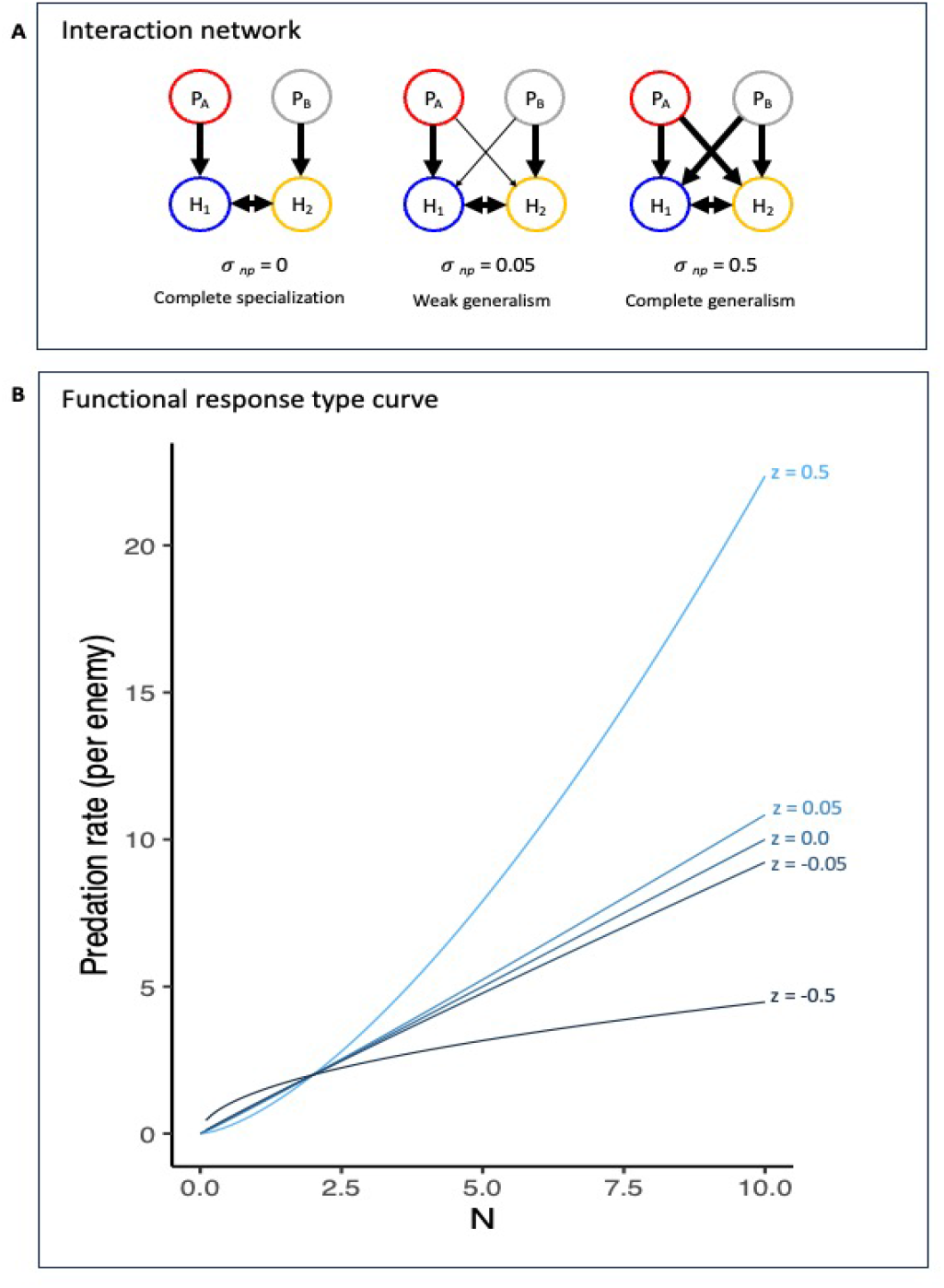
(A) Degree of specialization on the interaction network. The figure illustrates how changing pathogen specificity alters the structure of the model, arrows signify feeding preference (boldness of the arrow signifies the strength of preference). Complete specialization (left): the enemies have a strong preference and only feed on their preferred victim. Intermediate specificity (middle): the enemies show a slightly relaxed feeding preference, and supplement their feeding on their preferred victim with some feeding on non-preferred victim. The interaction between the enemy and their non-preferred victim is weak compared to the interaction between the enemy and their preferred victim. Complete generality (right): the enemies do not show any preference for one victim over the other. Victim species (*H*_1_ and *H*_2_) compete with themselves and with each other with equal magnitude, and are regulated only by natural enemies. Enemies do not compete with each other directly, but might compete indirectly through shared victim. (B) The curves illustrate degrees of functional response types. In our model *z* =0 signifies a linear response of enemies to their victim’s density, *z* < 0 3is3saturating, and *z* > 0 is an accelerating response.

### Scaling and parameter reduction

To facilitate analysis and interpretability, we performed a scaling and parameter reduction (see Appendix A1 for details). We considered new variables relative to arbitrary reference scales, i.e. 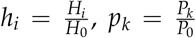, and 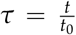, where we chose the references scales *H*_0_, *P*_0_, and *t*_0_ that make the scaled victim densities dimensionless (since they are measured relative to the victim carrying capacity when it is on its own). This operation also allowed us to reduce the total number of parameters. Using the following reference scales, and the reduced set of scaled parameters shown in Table 2

**Table 2:** Scaled parameters and their definition.

| Parameter | Definition |
| --- | --- |
| $\sigma_p \equiv H_0 s_p = s_p / \alpha$ | scaled susceptibility of victim species to enemy specialized on it |
| $\sigma_{np} \equiv H_0 s_{np} = s_{np} / \alpha$ | scaled susceptibility of victim species to enemy not specialized on it |
| $\phi \equiv f c t_0 = f c / r$ | scaled conversion of victim biomass to enemy biomass (or fecundity) |
| $\delta \equiv d t_0 = d / r$ | scaled intrinsic death rate of enemies |

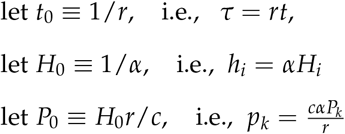

we transformed Eqs. 1 in terms of 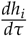, and 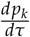, for *i* =1, 2 and *k* =*A, B*:

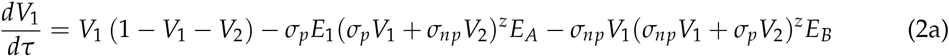

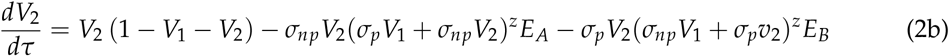

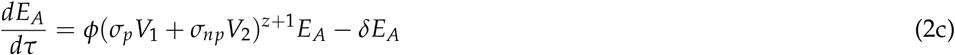

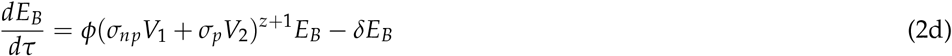

### Equilibria and feasibility

#### Feasibility

We found the coexistence equilibrium point (the one where all species can have a non-zero population size) of the model described by Eqs. 2a-d, and then determined the conditions for it to be feasible, i.e., for all densities to be positive. We then used these conditions to consider how the degree of specialization (i.e. the size of *σ*_*np*_), and other model parameters, impact feasibility.

#### Local Stability

We computed the Jacobian matrix associated with Eqs. 2a-d for equilibrium points with positive, non-zero densities for every species (i.e., feasible coexistence equilibria). The eigenvalues of the Jacobian were evaluated to assess the local stability of the coexistence equilibrium point (i.e. when the real part of the “dominant” eigenvalue (ReDEv)—the eigenvalue with the largest real part—is *<* 0). We considered the overall possibility for local stability analytically in the simpler *z* =0 case. To assess the effect of victim (or host) specificity on stability we calculated the eigenvalues of the Jacobian at the coexistence equilibrium while varying *σ*_*p*_ and *σ*_*np*_ between strict specificity (*σ*_*np*_ =0) and complete generality (*σ*_*np*_ =*σ*_*p*_). Relevant calculations can be found in the appendix A2.

#### Limit Cycles

At *z* =0, the system undergoes a Hopf bifurcation, with limit cycles emerging for *z* < 0. Using nonlinear dynamical systems techniques and Poincare theory, we identified these limit cycles and studied their stability. Specifically, if endogenous limit cycles exist, there must be a period (*t*) over which all four state variables return to their initial values by the end of the cycle. In other words, *h*_1_(0)= *h*_1_(*t*), *h*_2_(0) = *h*_2_(*t*), *p*_*A*_(0) = *p*_*A*_(*t*), *p*_*B*_(0) = *p*_*B*_(*t*). Using Newton’s method for numerical root finding, we solved for both this period *t* and the values of *h*_1_, *h*_2_, *p*_*A*_, and *p*_*B*_ at time = 0, given a reasonable initial guess (EcoEvo package, Parker and Chua 2012). Limit cycles identified using this approach may not be stable or unique. After a given limit cycle was detected, bifurcation continuation techniques were used to explore how variation in parameter values affected limit cycle properties across parameter space. To determine the stability of limit cycles, we calculated the corresponding set of Floquet exponents (EcoEvo package, Klausmeier 2008). When the real part of the dominant Floquet exponent (ReDFE) is greater than 0, the corresponding limit cycle is unstable. Otherwise, the ReDFE will be in the neighborhood of 0 (reflecting the neutral stability of perturbations that advance the system in the direction of the limit cycle), and the real part of the next-largest Floquet exponent provides a measure of the rate at which other small perturbations away from the limit cycle will decay (Strogatz 2018).

#### Numerical Simulation

We used numerical simulations to explore the behavior of the system away from the equilibrium point. We restricted our exploration to the feasible region of parameter space, within which we chose 100 random non-equilibrium starting population values for each combination of parameters and ran the numerical simulations for 5,000 time steps. Numerical simulations were performed using R 4.3.0 (R Core Team 2021). All R and Mathematica (Inc. 2021) code is available on GitHub (see Data Availability).

## Results

### Specialization and feasibility

This symmetric two-enemy, two-victim model has multiple fixed points, but only one coexistence equilibrium at:

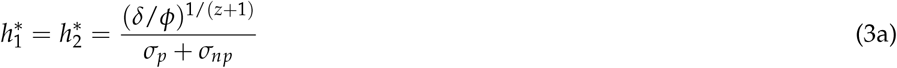

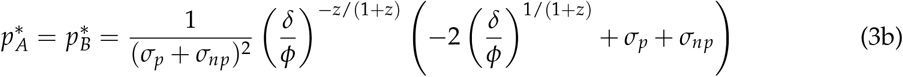

This equilibrium is feasible under all biologically relevant conditions where *δ, ϕ* are non-zero positive real values, *σ*_*p*_ and *σ*_*np*_ are positive real values, *z≠* −1, and

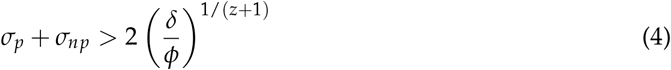

From equation (4), we can see that increasing the susceptibility of victims to both their specialist and non-specialist enemies contributes equally to satisfying the conditions for a feasible equilibrium. If enemies specialize completely on a single victim (i.e., *σ*_*np*_ = 0), *σ*_*p*_ has to be large enough on its own to exceed the right-hand side of the inequality in Eq. 4. If 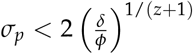, complete specialization leads to enemy extinction, and hence is incompatible with feasible four-species coexistence.

When the rate of scaled enemy mortality is lower than scaled enemy fecundity 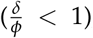, the right-hand side of the feasibility criteria shows a monotonic increase with *z* for *z* > −1 (the R.H.S. of the inequality is undefined at *z* = *−*1). Hence, as *z* increases so that the enemy functional response becomes less saturating (and eventually becomes accelerating once *z* > 0), lower values of *σ*_*p*_ and *σ*_*np*_ will be sufficient for the community equilibrium to be feasible.

### Local stability and numerical simulations when enemies have linear functional responses

Linear stability analysis demonstrated that the feasible equilibrium of the model when enemies have a linear functional response (*z* = 0) is not linearly stable. From the expressions for the eigenvalues of the Jacobian evaluated at the coexistence equilibrium for this case (below; see Appendix A2 for full derivation, and GitHub for Mathematica code), it can be determined that the real part of the dominant eigenvalue (ReDEv) of the coexistence equilibrium point is equal to 0. Specifically, the four eigenvalues of the Jacobian at the equilibrium point can be shown to take the form:

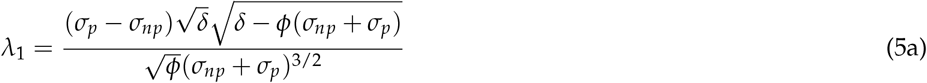

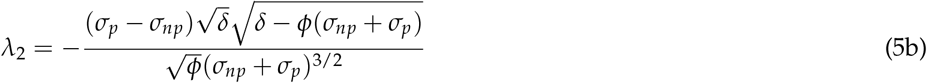

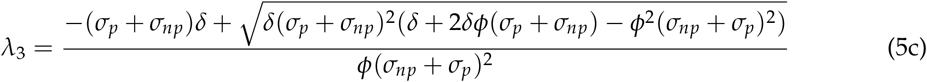

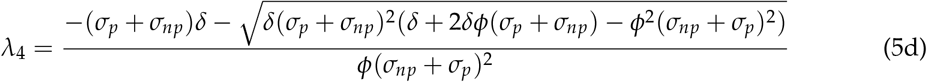

When the feasibility condition (Eq. 4) is met, the term under the square root in the numerator for eigenvalues *λ*_1_ and *λ*_2_ is always negative and hence these eigenvalues are purely imaginary (i.e. they have real parts = 0). It can also be shown that *λ*_3_ and *λ*_4_ have real parts ≤ 0 (Appendix A2). Using all this information together, we can conclude ReDEv = 0 for this case. Thus, the coexistence equilibrium point is *never* linearly stable (regardless of parameter values) in this case.

However, second and higher-order terms in the approximate dynamics of species populations near the equilibrium point may stabilize or destabilize the equilibrium. A true second-order analysis was beyond the scope of this study, but to explore this possibility we used numerical simulations, which indicated neutral cycles around the four-species coexistence equilibrium point (Fig. 2(A)). Specifically, we find all trajectories cycle around the equilibrium without converging on it. Trajectories started far from the equilibrium do get closer to it, but then stall out in a region around it, continuing to cycle there. However, trajectories do not seem to all approach a particular cycle, so there does not seem to be a stable limit cycle. The region of neutral cycles to which the system converges is smaller and tighter around the equilibrium when the enemy is more specialized (compare top and bottom rows of Fig 2(A)).

**Figure 2.**
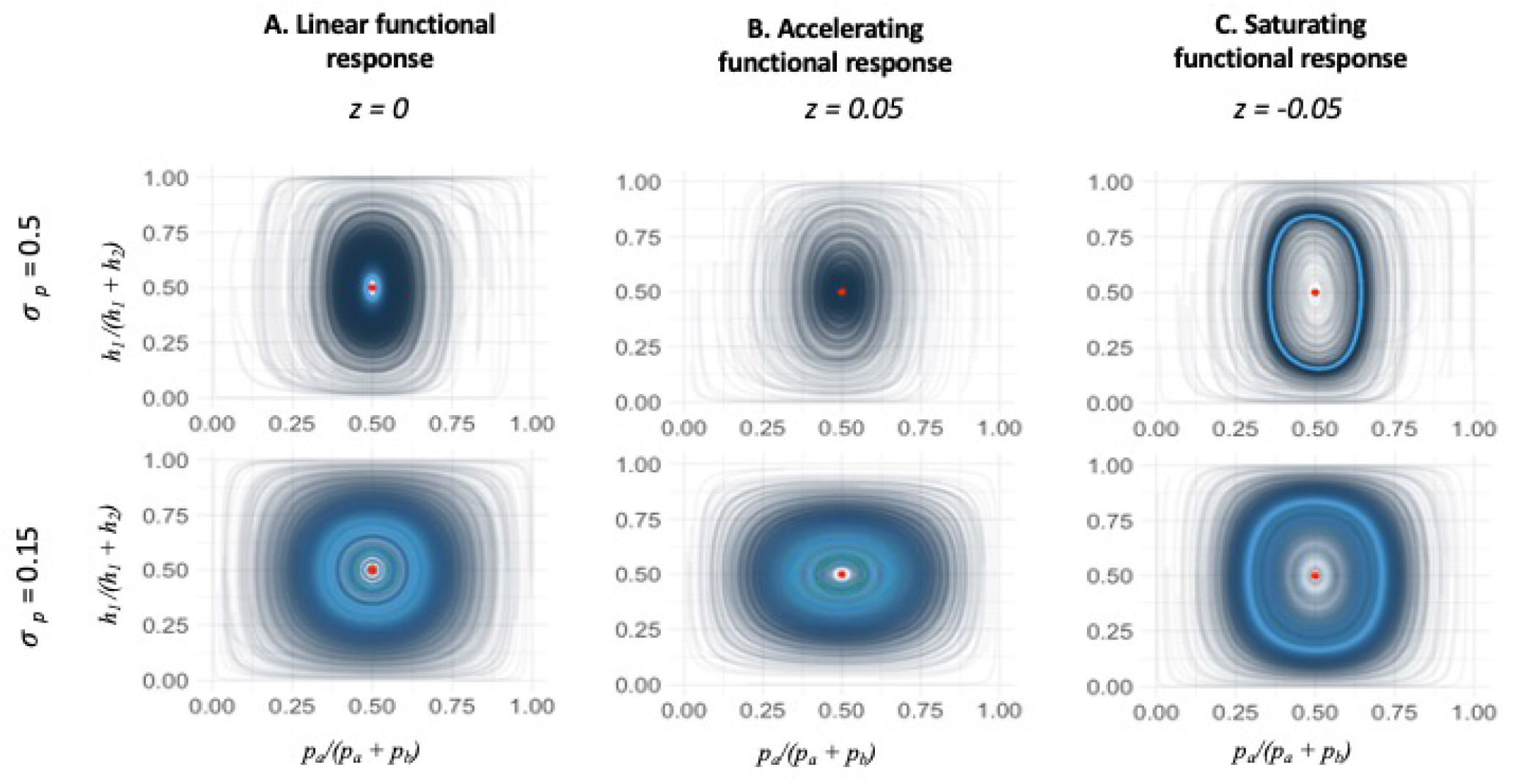
Numerical simulation showing trajectories of 100 randomly chosen non-equilibrium starting densities. The equilibrium point is denoted with a red dot in all the plots, and progression of time is denoted by the line color (dark to light blue as time progresses). The other parameters are defined as *σ*_*np*_ = 0.1, *δ* = 0.1, *ϕ* = 1. (A) *z* = 0, linear functional response. Enemy and victim population sizes cycle around the equilibrium point without converging all the way to it—they only converge to within a certain range of it, and appear to exhibit neutral cycles from there. The region in which the cycles appear neutral is larger when the difference between non-preferred (*σ*_*np*_) and preferred (*σ*_*p*_) susceptibility is small. (B) *z* = 0.05, accelerating functional response. Population sizes converge to the equilibrium point when non-preferred and preferred susceptibility have a higher difference (*σ*_*p*_ = 0.5 compared to *σ*_*p*_ = 0.15). (C) *z* = −0.05, saturating functional response. Trajectories converge to a limit cycle around the equilibrium. Here too, convergence happens faster (tighter light blue band) for *σ*_*p*_ = 0.5 compared to *σ*_*p*_ = 0.15.

### Local stability, numerical analysis and simulations when enemies have non-linear functional responses

We first summarize our results for non-linear functional responses and then provide greater detail below. We find that with equivalent competitor victims, only accelerating functional responses lead to stable equilibrium point behavior. Under saturating functional responses, the equilibrium point is locally unstable, but the potential for persistent coexistence through stable limit cycles and chaos remains, unless saturation is too strong, in which case the enemy populations cycle to extinction. See Fig. 3 for a summary of this behavior.

**Figure 3.**
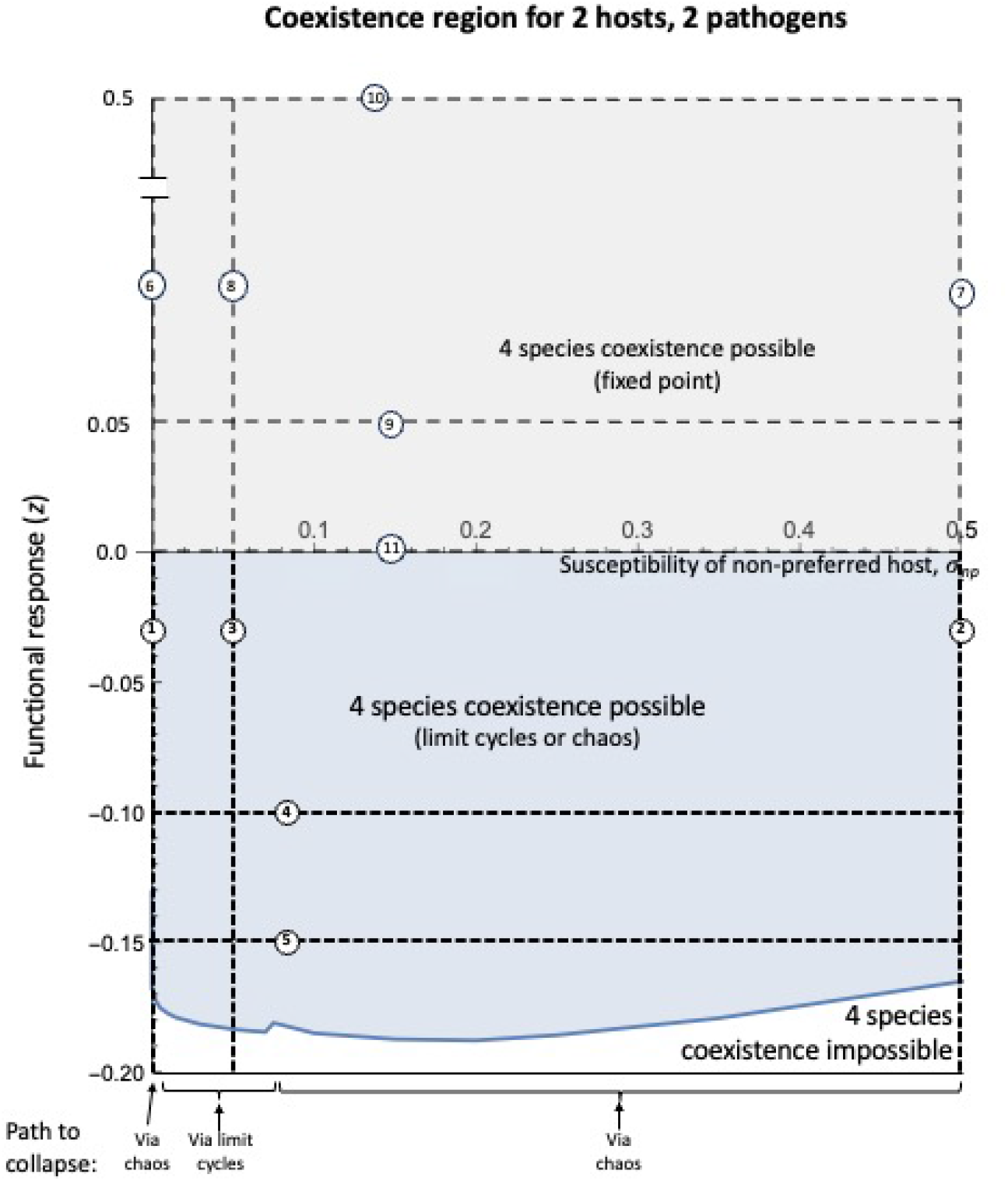
The region where the full, four species system exhibits persistence (either in the form of fixed points, one or more stable limit cycles, or chaotic attractors) depends on interactions between pathogen specificity (*σ*_*np*_) and the non-linearity of the functional response (*z*). Fixed point coexistence is shown in gray, and coexistence through limit cycles is shown in blue; outside of this region the system collapses into either a two-species system (one victim, one enemy) or extinction of all four species. Additional details on the stability and properties of dynamical outcomes appearing across slices through this parameter space (dashed lines, 1-11) are shown in Fig. 4 and Fig. 5 below.

Under an accelerating functional response, the stability of the equilibrium point typically increases (i.e. the real part of the dominant eigenvalue (ReDEv) becomes more negative and hence the equilibrium is approached more quickly) the greater the specificity (i.e. lower *σ*_*np*_) and the greater the degree of acceleration (*z*). However, note these can lower the equilibrium point below feasibility in the extreme. Under a saturating functional response, the stability of the limit cycle (i.e. how negative the real part of the Floquet exponent (ReDFE) is) is typically highest at intermediately high specificity and moderate degrees of saturation. These trends in dynamical stability are exhibited in Fig 2(B) and (C) which show numerical simulations for key cases, and in Figs. 4 and 5 which show the relevant measures of stability (ReDEv, ReDFE) for the slices through parameter space indicated in Fig. 3. See Appendix Figs. A3.1, A4.1-2, A5.1-3 for a more detailed look at stability measures and the dynamics through those slices.

**Figure 4.**
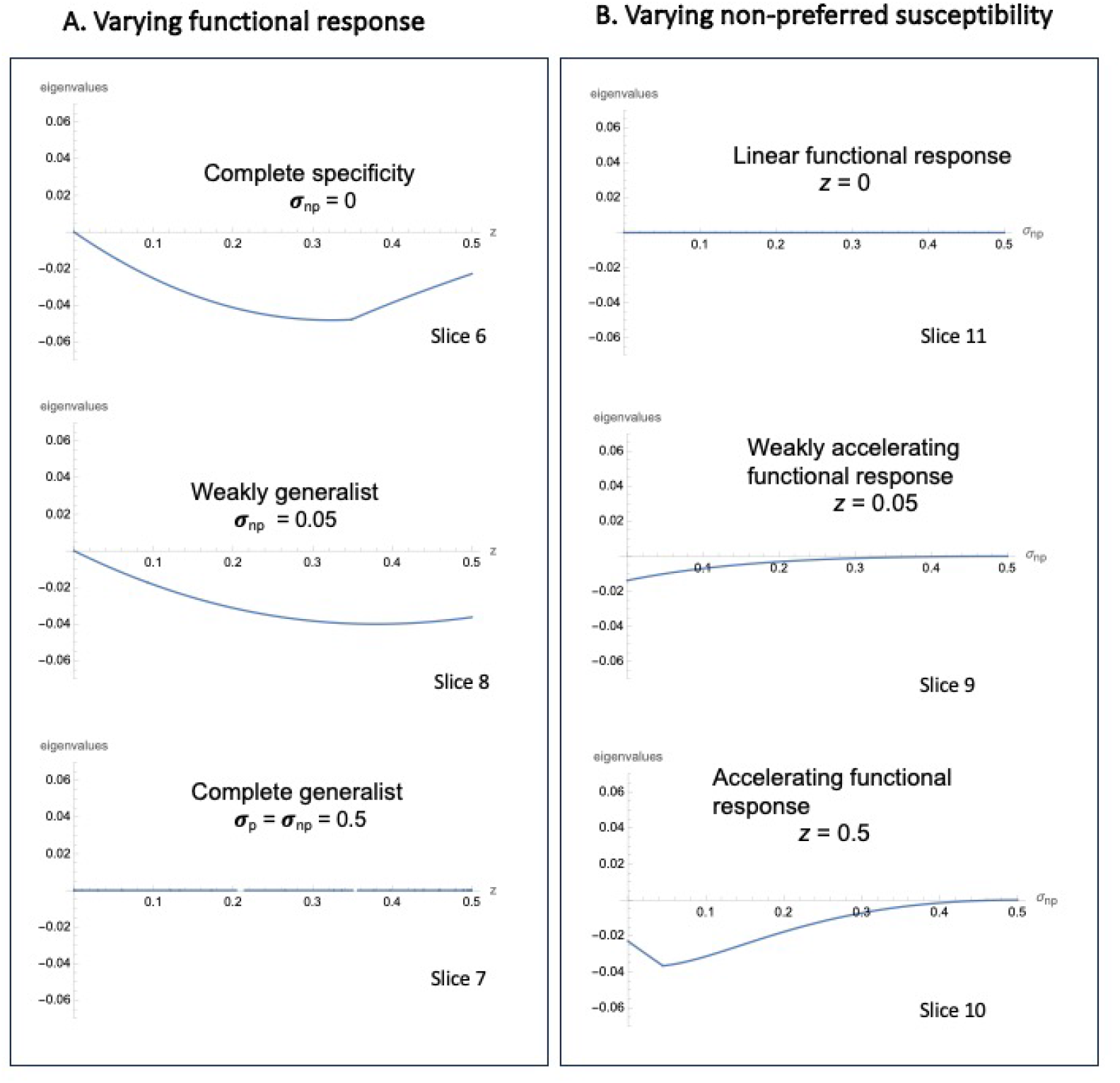
Effect of increasing the degree of acceleration with an accelerating functional response (varying *z ≥* 0) on the real part of the dominant eigenvalue (ReDEv) for three different levels of specificity *σ*_*np*_ (A), and impact of decreasing the degree of specialization *σ*_*np*_ for the linear (*z* = 0) and two degrees of accelerating (two *z >* 0) functional responses (B). The equilibrium point is *not* linearly stable or unstable (ReDEv = 0) under a linear functional response (*z* = 0). It is also *not* linearly stable or unstable for a complete generalist (bottom graph in (A), *σ*_*p*_ = *σ*_*np*_ = 0.5). When the enemies have some specificity, the stability typically increases (ReDEv gets more negative), as the functional response becomes more acclerating (top and middle graphs of (A)). Stability also typically declines with increasing generalization (higher *σ*_*np*_ a (middle and bottom graphs of (B)).

**Figure 5.**
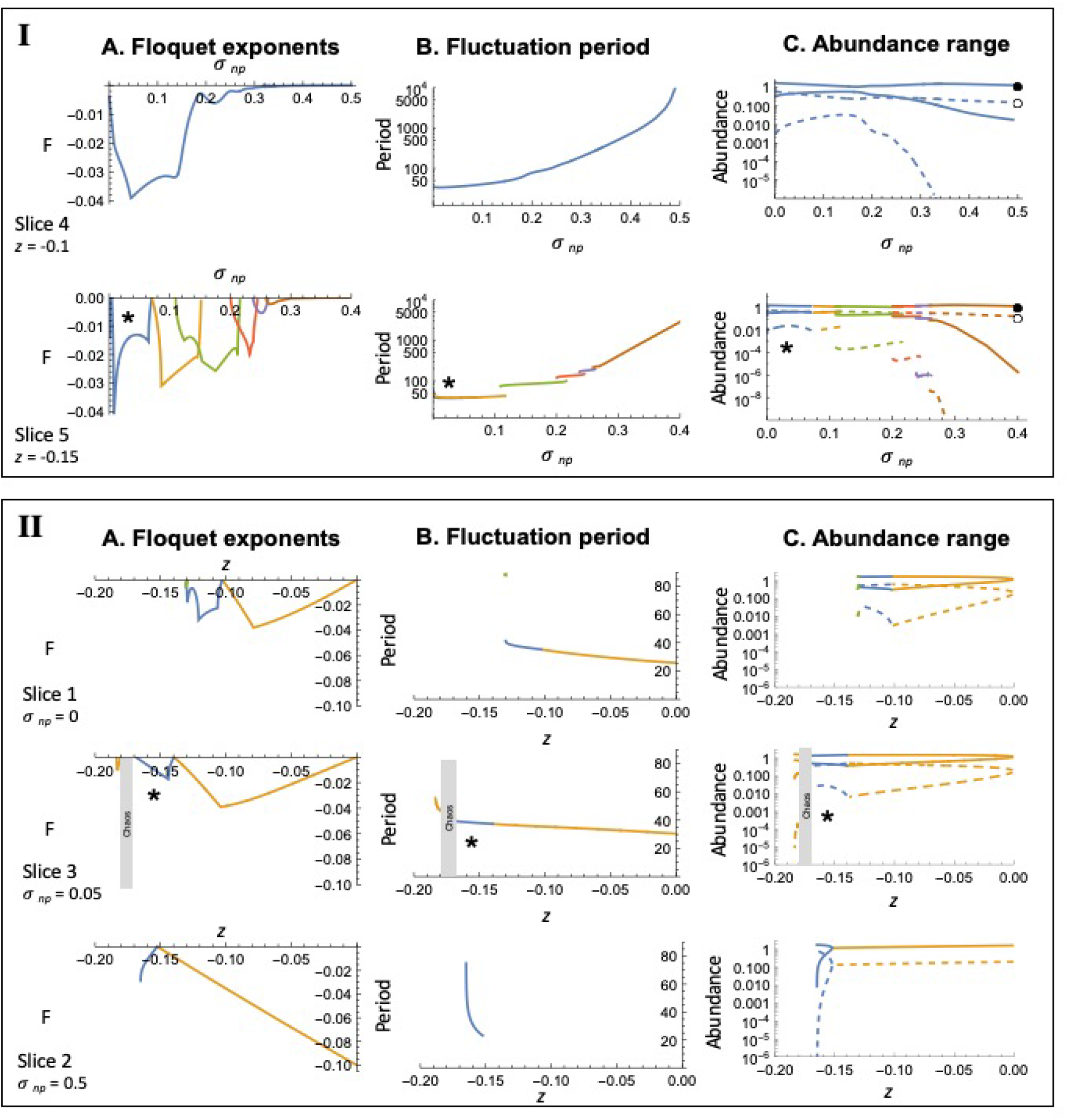
Real part of the dominant Floquet exponent (ReDFE), fluctuation period, and abundance range of a enemy-victim pair (*P*_*A*_ is solid, *H*_1_ is dashed) under limit cycles or chaos in the saturating functional response regime. Only negative values corresponding to stable limit cycles are shown. Different colors correspond to distinct types of limit cycles (orange, blue, green). Outcomes marked with a (*\**) correspond to a region of parameter space where for clarity only one of a pair of a set of alternative, asymmetric limit cycles is shown (see main text and Appendix A5 for details). (I) Effects of varying the degree of pathogen specialization (feeding on non-preferred victims, *σ*_*np*_) for parameter slices *z* = −0.1 (top row) and *z* = −0.15 (bottom row). Generally, as *σ*_*np*_ converges on *σ*_*p*_ (= 0.5) (feeding becomes completely general), the period of oscillations increases dramatically (note log scale in column B), the stability of the limit cycles becomes increasingly weak (ReDFE converges on 0), and victims drop to extremely low abundances at the lowest point of their cycle. In the bottom row, six distinct types of stable limit cycle are shown. Finally, as *σ*_*np*_ converges on *σ*_*p*_ (= 0.5), we generally see dramatic lengthening of cycles, where relatively abrupt changes in dominance interrupt long periods of nearly stationary dynamics. The peak abundance of victim and enemies across these cycles converges on the stable, fixed-point abundances of victim and enemy in the two species system (shown as the empty and filled black circles, respectively, right column). (II) Effects of varying the strength of the saturating functional response, *z*, along parameter slices *σ*_*np*_ = 0 (top row), *σ*_*np*_ = 0.05 (middle row), and *σ*_*np*_ = 0.5 (bottom row). Fluctuation period (middle column) generally increases as *z* decreases, and cannot be computed for chaotic attractors (middle row, gray boxes). Finally, the range of abundances generally increases as *z* decreases, with victim populations in particular declining to low abundances at low *z*. In the complete generality case (bottom row, *σ*_*np*_ = *σ*_*p*_ = 0.5), victims pairs and enemy pairs are effectively neutral with respect to each other, so the results shown are for the special case of the interaction of one enemy and one victim, instead of the 4 species case.

#### Accelerating functional response

The coexistence equilibrium point of this two-victim and two-enemy system is linearly stable when enemies have an accelerating response (*z >* 0) to victim densities and enemies are not complete generalists. Specifically, ReDEv *<* 0 for all *z >* 0 and *σ*_*np*_ *< σ*_*p*_ (see Fig. 4(A) and (B) and Appendix Fig. A3.1(A) and (B)). We also find that the degree of stability (how negative ReDEv is) tends to increase with the degree of acceleration of functional response (with *z*) (See Fig. 4(A) and compare Appendix Fig. A3.1(A) with A3.1(B)), except for higher values of *z* and complete specificity *σ*_*np*_ = 0 (right end of top graph in Fig. 4(A)). Furthermore, we see in Fig. 4(B) that stability tends to decline (ReDEv becomes less negative) with increasing generalization (increasing *σ*_*np*_), except for at low values of generalization (small *σ*_*np*_) with a quickly accelerating functional response (*z* = 0.5) (left end of bottom graph in Fig. 4(B)). Numerical simulations starting at random non-equilibrium values show that trajectories do indeed go to the equilibrium point from a wide range of starting values (the stability is not just local stability). Furthermore, they converge on the equilibrium point very quickly when enemies have a high baseline specificity to their victims (see upper row in Fig. 2(B), for which *σ*_*p*_ is high and *σ*_*np*_ is low, and hence specificity is high). For a complete generalist, this equilibrium point stability no longer occurs. Instead, the populations converge toward the equilibrium somewhat but then appear to neutrally cycle around the equilibrium point (see bottom row in Fig. 2(B)). See Appendix Fig A4.1 and A4.2 for an additional closer look at population dynamics across degree of acceleration *z* and degree of specificity (along the parameter slices indicated in Fig. 3).

#### Saturating functional response

Our numerical simulation (Fig. 2(C)) with random non-equilibrium starting values show that trajectories convergence on cycles (light blue band) around the equilibrium point (red dot). Moreover, they converge on the limit cycles much faster when enemies have a high baseline specificity to their victims (see upper row of Fig. 2(C) for which *σ*_*p*_ is high and *σ*_*np*_ is low). Below we present the results of further analysis of the dynamics using Floquet theory.

In the saturating functional response cases (*z <* 0), the equilibrium point is linearly *unstable*. Specifically, the real part of the dominant eigenvalue (ReDEv) is positive across all values of susceptibilities when functional responses are saturating (see Appendix Fig. A3.1(D)and(E)). However, simulations and Floquet exponent analysis indicate that the dynamics away from the equilibrium point are complex, and can involve species persisting through stable limit cycle or chaos, unless the degree of saturation is large (i.e. *z* more negative), in which populations go extinct even if the equilibrium point is feasible (see Fig. 3). The stability of limit cycles, i.e. how negative the real part of the Floquet exponent is, is greatest at intermediately high specificity and moderate degrees of saturation (see Fig. 5 for the Floquet exponent, fluctuation period, and abundance range of limit cycles across parameter slices indicated in Fig. 3 Appendix Fig. 5.1-3 for a numerical simulations of population dynamics along those parameter slices, Fig. 2(C) for numerical simulations for key cases).

Within the coexistence region, the four-species community exhibits a complicated array of dynamical features, including both diverse limit cycles and chaos depending on *z* and *σ*_*np*_. The boundary between coexistence and collapse of the four species community occurs as either limit cycles or chaotic oscillations become unstable, depending on the difference between the values of *σ*_*np*_ and *σ*_*p*_. Decreases in the difference between *σ*_*np*_ and *σ*_*p*_ (i.e. increased generality) are associated with increases in the Floquet exponent (i.e., decreased stability), except at very low *σ*_*np*_ (Fig. 5 I). In this sense, the stability of the limit cycle is highest at intermediately high specificity. As the enemies relax their preference (*σ*_*np*_ approaches *σ*_*p*_), the decrease in stability is accompanied by a sharp increase in period length and minimum abundance. As *z* decreases (saturation of the enemies emerges at lower victim densities), the general relationships between specificity and stability of the system remain (Fig. 5 I, bottom row). However, instead of all of the solutions emerging from continuous changes in one class of limit cycles across the range of *σ*_*np*_, we detected the existence of (at least) six distinctly different stable limit cycles, sometimes occurring for the same value of *σ*_*np*_ (Fig. 5 I, bottom row). See Appendix Fig.A5.2-3 for an additional closer look at the limit cycles as *σ*_*np*_ varies. As the use of the alternative victim increases (around *σ*_*np*_ ≈ 0.3 for the current set of parameters in Fig. 5 I, bottom row) the periods get very large, and abundances show massive fluctuations spanning more than five orders of magnitude. One enemy-victim pair is dominant at any given time, and step-wise replacement of the dominant pair by the non-dominant takes a very long time. This gives way to the two-species model via an infinite-period bifurcation (Hsu et al. 2009). Additional details are shown in Appendix A5.

In the case of complete specialization, in which case enemy dynamics are only connected through competition between their respective victims, we see three distinct limit cycles unfold across the *z*-range (Fig. 5 II, top row). As *z* decreases, an initial limit cycle loses stability, giving rise to a pair of alternative, asymmetric limit cycles (see Appendix A4). These in turn lose stability at yet lower values of *z*, replaced briefly by cycles with a doubled period length, before four-species coexistence becomes impossible as *z* decreases further. At this point, a stable, fixed coexistence point emerges for a single enemy-victim pair, which becomes oscillatory, and then itself collapses.

Even a small amount of non-preferential feeding by the enemies (0 < *σ*_*np*_ ≪ *σ*_*p*_) expands the range of *z* consistent with four-species coexistence and decreases Floquet exponents markedly. As *σ*_*np*_ increases further however, the Floquet exponents increase and the period of the limit cycles increase, both indicating reduced stability and longer return times when the system is perturbed. When enemies do not distinguish between victims at all (complete generality, *σ*_*np*_ = *σ*_*p*_), the dynamics of pairs of both enemies and victims are effectively neutral with respect to each other and period length increases rapidly towards infinity (line (2) from Fig. 3). This coincides with the gradual collapse of the limit cycle solution (via an infinite period bifurcation (Strogatz 2000, p. 262), into the fixed-point equilibrium exhibited by the simpler, two-species system with one enemy and one victim. This two-species system exhibits stable fixed point solutions for a wide range of *z <* 0, which lose stability and generate limit cycles for more extreme (lower) values of *z* before finally collapsing.

## Discussion

Prior work on the influence of enemies on host coexistence has found that more specialized enemies generate more coexistence potential (Sedio and Ostling 2013, Chesson 2018). However, this work has largely ignored enemy dynamics and the potential for enemy extinction. Where it has included enemy dynamics, it has assumed the enemy has additional unmodeled hosts or a constant rate of immigration into the community, and has focused only on a simple linear functional response. Here we studied a model of two competitively-equivalent victim species regulated by two enemy species and showed that the feasibility criteria for coexistence are easier to attain as enemies become less specialized. The effect of enemy specificity on stability is more complex and depends on the functional responses of the enemies, which are in themselves also crucial to the outcome of the system’s dynamics. For the enemies to generate stable equilibrium point co-existence of the victims requires an accelerating functional response of enemies to victim density, which most studies suggest are rarely encountered in nature (Dunn and Hovel 2020; Hopkins et al. 2020), but observations for the fungal pathogens that may play a strong role in tree species coexistence suggest otherwise (Bagchi et al. 2010). Linear functional responses (Type 1) led to neutral stability, and when the functional response of the enemy to victim densities is saturating, complex dynamics involving stable limit cycles, chaos, and extinction can result. Higher specificity tends to increase stability (more negative dominant real part of the Floquet exponent) when there is stable equilibrium point behavior, but the stability of limit cycle behavior is promoted by higher, but not complete specificity. Stronger saturating functional responses initially promoted stable coexistence before generating ever more extreme cycles, chaos and collapsing. Our results highlight the importance of the consideration of enemy dynamics in the growing body of theory of enemy-mediated coexistence.

Our model indicates that increasing victim specificity can increase the stability of a two-victim, two-enemy coexistence equilibrium under certain conditions, a result that coincides with previous work (Adler and Muller-Landau 2005; Bruijning et al. 2023; Chesson 2018; Kirchner and Roy 2002; Sedio and Ostling 2013). However, by explicitly modeling enemy population dynamics within a closed community, our model revealed limits on the capacity of enemies to enable coexistence of their victims: specifically, the vulnerability of enemies to extinction themselves. This limit has important implications. Hypothetically, specialized enemies could regulate their victims’ populations and prevent competitive exclusion while, simultaneously, avoiding competition with each other by specializing on different victim species. This self-reinforcing cycle could potentially support limitless diversity. However, species in higher trophic ranks are more susceptible to extinction, a characteristic reinforced by strict specialization (Grenfell et al. 1995; Grenfell and Harwood 1997; Holt et al. 1999; Thrall et al. 2003; De Castro and Bolker 2004; Gravel et al. 2011; Seabloom et al. 2015). This high extinction risk will likely be exacerbated in very diverse communities, where enemies are dependent on victims that occur at low density (May 1991; Woolhouse 2001). Furthermore, if victim populations are primarily regulated by one species of enemy, enemy extinction would release some victim species from population regulation and possibly lead them to competitively exclude other victim species. Such extinction of highly specialized enemies would be unsurprising and might act to limit an otherwise infinite accumulation of specialized victim-enemy pairs. Our current model did not account for demographic and/or environmental stochasticity but it should be noted that they could further destabilize a community with many highly specialized victim-enemy pairs (Jeltsch et al. 2019). Overall, our analyses in this study suggest that while increasing host specificity might aid species coexistence, the relationship between specificity and diversity is unlikely to be monotonically positive.

Host specificity is a function of several different biological processes, and most models approximate those processes with abstract parameters that only incompletely reflect natural systems (Antonovics 2017; McCallum et al. 2017; Webster et al. 2017). In keeping with this trend, our definition of specificity relied solely on the scaled susceptibilities of preferred and non-preferred victim species to the enemy species. Our model held enemy fecundity and mortality constant and independent of which victim species were consumed. Although these simplifications increase model tractability, they hinder empirically-based parameterizations of models. A more mechanistic representation of victim-enemy interactions could allow the component enemy traits (e.g., fecundity and mortality) to depend on victim species identity (e.g., as would be expected in host-pathogen scenarios). Such models could also represent specialization through variation among species in the component biological processes of host competence, virulence, defense responses, dispersal, and the trade-offs between them (Thrall 2003; Messinger and Ostling 2009; Lion and Boots 2010; Messinger and Ostling 2013). These quantities can be empirically measured, allowing for an assessment of not only whether certain parameter values are theoretically compatible with species coexistence, but also whether naturally occurring traits are consistent with promoting species coexistence. Such links between empirical measurements and theory present opportunities for mechanistic theories of species coexistence and diversity, but are challenging to achieve in practice.

The functional response of enemies to victim densities (*z* in our models) was the second feature critical to a four-species coexistence. Like specificity, functional responses also reflect multiple aspects of the ecology of both enemies and victims, including spatial dynamics, physiology, habitat structural complexity, enemy learning, victim rarity, as well as environmental factors like temperature (Dunn and Hovel 2020; Pascual et al. 2011). The majority of enemy-victim models assume a linear relationship (type I functional response) between victim density and attack rates by enemies (e.g, Lotka-Volterra models (Lotka 1925, Volterra 1927; SI models Kermack and McKendrick 1927; Hopkins et al. 2020). However, type I functional responses are rare in natural enemy-victim interactions (McCallum et al. 2001; Orlofske et al. 2018; Hopkins et al. 2020) and are generally limited to filter feeders (Jeschke et al. 2004). Reviews and meta-analyses of the prevalence of functional responses have generally found Type 2, followed by Type 3, functional responses more common in nature, including in carnivorous (Dunn and Hovel 2020) and pathogenic systems (Hopkins et al. 2020). In the model presented here, coexistence of all four species was feasible under certain conditions with a linear functional response (*z* = 0), but the equilibrium was neutrally stable (i.e., the system did not return to the equilibrium point if perturbed).

Stable coexistence in the feasible region at an equilibrium point was only observed with an accelerating relationship between attack rate by enemies and victim density (*z >* 0). However, empirical examples of such accelerating relationships appear rare in nature (but see an example of rapid overcompensating density-dependent mortality in seedlings driven by fungal pathogens in Bagchi et al. 2010), although that might partly reflect a paucity of studies designed to identify it. At some point, an accelerating functional response must saturate; however, accelerating functional responses might best describe dynamics at low victim densities or at restricted spatio-temporal scales. Indeed, the original formulation of the well-known Janzen-Connell hypothesis (Connell 1971; Janzen 1970) implicitly invokes an accelerating functional response of enemies at local scales by proposing decreased seedling recruitment when conspecific seed density is high (Bagchi et al. 2010). An accelerating functional response to victim densities might occur when enemy dynamics are rapid (e.g., many fungal pathogens) and dispersal is local (e.g., splash and contact dispersing pathogens). Our model suggests that under this assumption of an accelerating functional response, coexistence is stabilized, even when enemies are moderately polyphagous. While the stability of coexistence increases with specialization, eventually enemies become too specialized to maintain positive population growth when their preferred victim goes through low-density periods, at which point coexistence collapses. Although previous studies have demonstrated that non-specialized enemies can maintain victim coexistence (Chesson 2018; Sedio and Ostling 2013), our analyses suggest this is true even within a closed enemy population (i.e., not supported by immigration or alternative victim species).

Saturating functional responses are the most commonly observed response type in nature. Our analysis reveals the complicated array of dynamics behind both coexistence and collapse of a two-enemy-two-victim system under saturating functional responses. These dynamical features, including limit cycles, chaotic attractors, and heteroclinic cycles were also detected by Hsu et al. (2009), who studied a similar model. They examined a two-enemy-two-victim system, with saturating functional responses, but assumed complete specialization and focused on the effects of nutrient enrichment on coexistence and stability. Complementing their results, we show that even though the coexistence equilibrium point is unstable, the four-species system can be persistent, depending on interactions between enemy and victim’s non-preferred susceptibility (*σ*_*np*_) and the degree of the saturating functional response (*z*).

For highly specialized enemies (*σ*_*np*_ *≪ σ*_*p*_), limit cycles lost stability and gave way to chaotic oscillations. Chaotic solutions extended until *σ*_*np*_ values were very nearly zero, at which point the four species system collapsed as one victim was driven to extinction. This indicates that even a small amount of non-preferential feeding is able to rescue and stabilize the system which would otherwise collapse when enemies are completely specialized on their victim. Furthermore, as enemies relaxed their feeding preference further such that *σ*_*np*_ approaches *σ*_*p*_, we observed a dramatic increase in the period length of fluctuation of the limit cycles, a characteristic of infinite period bifurcations (Strogatz 2000) (see Appendix for details). As a result of these very slow oscillations, both enemy and victim reach extraordinarily low densities with one enemy-victim pair dominating at any given time. The community essentially behaves like a one-enemy-victim system as step-wise replacement of the dominant pair by the non-dominant takes a very long time. In summary, the four species system persists over higher degrees of saturation (lower values of *z*) when enemies show intermediate specificity. Biologically, it means that some degree of polyphagy can in fact mitigate the risk of enemy extinction, and a community of two enemies and two victims can tolerate a higher degree of saturating functional responses and coexist when enemies relax their feeding preference.

Further work with alternative and in particular more realistic model formulations will be necessary to determine whether this result is a general pattern in enemy-victim systems. Studying type III functional responses (combining accelerating and saturating parts) (Uszko et al. 2015), or better yet explicitly modeling the functional response as arising from spatial and/or behavioural interactions, could shed light on the mechanisms at play. Our current model is deterministic in nature, where both enemy and victim populations reach very low densities but do not go extinct. Stochastic noise of any kind will potentially affect the system dramatically given low densities (Jeltsch et al. 2019) and even more so in closed habitats of finite sizes (Schreiber et al. 2020). Additionally, extinction risk has been shown to increase or decrease depending on the probability of extreme events in the case of positively correlated stochastic fluctuations (Schwager et al. 2006). We believe that a consideration of demographic and environmental stochasticity will be crucial to our understanding of the mechanistic nature of enemy-mediated victim coexistence, and will assist us in establishing a well-rounded understanding of the complexities of enemy-victim interactions. These more realistic model formulations can then be used to reconsider key questions about enemy-mediated coexistence that are under debate, such as whether it is a mechanism that is robust to differences in species intrinsic demographic performance (Chisholm and Fung 2020; Smith 2022).

## Conclusion

Specificity of enemy-victim interactions has long been cited as essential to enemy-mediated co-existence, and previous studies (Sedio and Ostling 2013, Chesson 2018) have suggested that increased specificity supports coexistence. However, in this paper we show that the effect of specificity is complex when enemy population dynamics are explicitly modelled in a closed system. Even though victim coexistence is supported by an increase in specificity, it comes at the cost of a smaller parameter space over which the enemy populations are feasible (non-zero population density), and hence poses a higher risk of enemy extinction. Additionally, the functional response of the enemies is pivotal in determining whether coexistence of all enemy and victim species is possible. We found that a two-enemy and two-victim system stably coexists under an accelerating relationship between enemy attack and victim density, a result that is surprising at first; however, accelerating functional responses might best describe dynamics at low victim densities or at restricted spatio-temporal scales. The functional response of enemies to victim density moderates the relationship between specificity and the stability of coexistence. When enemies exhibit saturating functional response, the system’s equilibrium coexistence is unstable, but the four-species system persists as stable limit cycles or chaotic attractors. We showed that some degree of polyphagy mitigates the risk of enemy extinction and makes the community persist over a greater range of saturating functional responses. The nature of most functional responses between enemies and their victims is saturating, and our study identifies potential mechanisms of coexistence without the requirement of strict specificity of enemy-victim interactions.

