## Supplementary material for "Influence of natural enemy specificity and functional response on victim coexistence"

### Appendix

In our model, two enemy species $(P_A, P_B)$ consume two victim species $(H_1, H_2)$, and the encounter leads to losses and gains in biomass of the victim and enemy species respectively:

$$\frac{dH_1}{dt} = H_1 r\left(1 - \alpha H_1 - \alpha H_2\right) - c s_p H_1 (s_p H_1 + s_{np} H_2)^z P_A - c s_{np} H_1 (s_{np} H_1 + s_p H_2)^z P_B \tag{1a}$$
$$\frac{dH_2}{dt} = H_2 r\left(1 - \alpha H_1 - \alpha H_2\right) - c s_{np} H_2 (s_p H_1 + s_{np} H_2)^z P_A - c s_p H_2 (s_{np} H_1 + s_p H_2)^z P_B \tag{1b}$$
$$\frac{dP_A}{dt} = fc(s_p H_1 + s_{np} H_2)^{z+1} P_A - dP_A \tag{1c}$$
$$\frac{dP_B}{dt} = fc(s_{np} H_1 + s_p H_2)^{z+1} P_B - dP_B \tag{1d}$$

### Appendix A1: Parameter and dimension reduction

We started with a fully symmetric system with two-enemies and two-victims (Eqs 1a-d). The contact between enemies and victims are assumed to follow a power-law function of the victims that enemies consumes, weighted by the preference of that enemy has for each victim, i.e., for $H_1$, we have $(s_p H_1 + s_{np} H_2)^z$ for enemy $A$; $(s_{np} H_1 + s_p H_2)^z$ for enemy $B$. We chose new variables relative to arbitrary scales $h_i = \frac{H_i}{H_0}$, $p_k = \frac{P_k}{P_0}$, and $\tau = \frac{t}{t_0}$. We then chose reference scales $H_0, P_0, t_0$ to make the scaled host densities dimensionless.

We assumed $t_0 \equiv 1/r$, i.e., $\tau = rt$; $H_0 \equiv 1/\alpha$, i.e., $h_i = \alpha H_i$; $P_0 \equiv H_0 r/c$, i.e., $p_k = \frac{c\alpha P_k}{r}$

We could then reduce the number of parameters $h_i = \alpha H_i$, $\sigma_p = \frac{s_p}{\alpha}$, $\sigma_{np} = \frac{s_{np}}{\alpha}$, $p_k = \frac{c\alpha}{r} P_k$, $\phi = \frac{fc}{r}$, $\delta = \frac{d}{r}$, and $\tau = rt$, and rewriting the original system in terms of the redefined parameters for $H_i, (i = 1)$ as

$$\begin{aligned}
\frac{dh_1}{d\tau} &= \frac{\alpha}{r}\frac{H_1}{dt} \\
&= \frac{\alpha}{r}\left(\frac{r}{\alpha}\alpha H_1\left(1 - \alpha H_1 - \alpha H_2\right) - \frac{s_p}{\alpha}\alpha H_1\left(\frac{s_p}{\alpha}\alpha H_1 + \frac{s_{np}}{\alpha}\alpha H_2\right)^z cP_A - \frac{s_{np}}{\alpha}\alpha H_1\left(\frac{s_{np}}{\alpha}\alpha H_1 + \frac{s_p}{\alpha}\alpha H_2\right)^z cP_B\right) \\
&= \frac{\alpha}{r}\frac{r}{\alpha} h_1\left(1 - h_1 - h_2\right) - \sigma_p h_1 (\sigma_p h_1 + \sigma_{np} h_2)^z \frac{c\alpha}{r} P_A - \sigma_{np} h_1 (\sigma_{np} h_1 + \sigma_p h_2)^z \frac{c\alpha}{r} P_B \\
&= h_1(1 - h_1 - h_2) - \sigma_p h_1 (\sigma_p h_1 + \sigma_{np} h_2)^z p_A - \sigma_{np} h_1 (\sigma_{np} h_1 + \sigma_p h_2)^z p_B
\end{aligned} \tag{2}$$

Similarly for victim 2, we get $\frac{dh_2}{d\tau} = h_2(1 - h_1 - h_2) - \sigma_{np} h_2 (\sigma_{np} h_1 + \sigma_p h_2) p_A - \sigma_p h_2 (\sigma_{np} h_1 + \sigma_p h_2) p_B$.

For enemy $A$, we can write

$$\begin{aligned}
\frac{dP_A}{d\tau} &= \frac{\frac{c\alpha}{r}}{r}\frac{dP_A}{dt} \\
&= \frac{\frac{c\alpha}{r}}{r}\left(f\left(\frac{s_p}{\alpha}\alpha H_1 + \frac{s_{np}}{\alpha}\alpha H_2\right)^{z+1} cP_A - dP_A\right) \\
&= \frac{fc}{r}(\sigma_p h_1 + \sigma_{np} h_2)^{z+1}\frac{c\alpha}{r} P_A - \frac{d}{r}\frac{c\alpha}{r} P_A
\end{aligned}$$

$$\frac{dp_B}{d\tau} = \phi(\sigma_p h_1 + \sigma_{np} h_2)^{z+1} p_A - \delta p_A \tag{3}$$

And similarly for enemy $B$ we get $\frac{dp_B}{d\tau} = \phi(\sigma_{np} h_1 + \sigma_p h_2)^{z+1} p_B - \delta p_B$

The full community of 2-enemy-2-victim can now be written as

$$\frac{dh_1}{d\tau} = h_1\left(1 - h_1 - h_2\right) - \sigma_p h_1(\sigma_p h_1 + \sigma_{np} h_2)^z p_A - \sigma_{np} h_1(\sigma_{np} h_1 + \sigma_p h_2)^z p_B \tag{4a}$$

$$\frac{dh_2}{d\tau} = h_2\left(1 - h_1 - h_2\right) - \sigma_{np} h_2(\sigma_p h_1 + \sigma_{np} h_2)^z p_A - \sigma_p h_2(\sigma_{np} h_1 + \sigma_p h_2)^z p_B \tag{4b}$$

$$\frac{dp_A}{d\tau} = \phi(\sigma_p h_1 + \sigma_{np} h_2)^{z+1} p_A - \delta p_A \tag{4c}$$

$$\frac{dp_B}{d\tau} = \phi(\sigma_{np} h_1 + \sigma_p h_2)^{z+1} p_B - \delta P_B \tag{4d}$$

It is worth noting that the scaled victim densities $h_i = \alpha H_i$ are the victim density relative to the victim carrying capacity when it is on its own $(1/alpha)$ and hence are dimensionless, and that the scaled enemy densities $p_k = \frac{c\alpha}{r} P_k$ consider the enemy density relative to the host carrying capacity density when it is on its own $(1/\alpha)$, scaled by (i.e. divided by) the relative rates of host intrinsic growth and infection $(r/c)$.

The coexistence equilibrium of the reduced model is at

$$h_1^* = h_2^* = \frac{(\delta/\phi)^{1/(z+1)}}{\sigma_p + \sigma_{np}} \tag{5a}$$

$$p_A^* = p_B^* = \frac{1}{(\sigma_p + \sigma_{np})^2}\left(\frac{\delta}{\phi}\right)^{-z/(1+z)}\left(-2\left(\frac{\delta}{\phi}\right)^{1/(1+z)} + \sigma_p + \sigma_{np}\right) \tag{5b}$$

And this equilibrium is feasible under all biologically relevant conditions where $\delta, \phi$ are non-zero positive real values, $\sigma_p$ and $\sigma_{np}$ are positive real values, $z \neq -1$, and

$$\sigma_p + \sigma_{np} > 2\left(\frac{\delta}{\phi}\right)^{1/(z+1)} \tag{6}$$

The Jacobian of the reduced model(Eq (4)) evaluated at the coexistence equilibrium is

$$\begin{pmatrix}
\frac{-(\sigma_p+\sigma_{np})(\sigma_p^2+\sigma_{np}^2)z+(\frac{\delta}{\phi})^{(\frac{1}{1+z})}(-(\sigma_p+\sigma_{np})^2+2(\sigma_p^2+\sigma_{np}^2)z)}{(\sigma_p+\sigma_{np})^3} & -\frac{2\sigma_p\sigma_{np}(\sigma_p+\sigma_{np})z+(\frac{\delta}{\phi})^{(\frac{1}{1+z})}((\sigma_p+\sigma_{np})^2-4\sigma_p\sigma_{np}z)}{} & -\frac{\left((\frac{\delta}{\phi})^{\frac{1}{1+z}}\right)^{1+z}\sigma_p}{\sigma_p+\sigma_{np}} & -\frac{\left((\frac{\delta}{\phi})^{\frac{1}{1+z}}\right)^{1+z}\sigma_{np}}{\sigma_p+\sigma_{np}} \\
-\frac{2\sigma_p\sigma_{np}(\sigma_p+\sigma_{np})z+(\frac{\delta}{\phi})^{(\frac{1}{1+z})}((\sigma_p+\sigma_{np})^2-4\sigma_p\sigma_{np}z)}{} & \frac{-(\sigma_p+\sigma_{np})(\sigma_p^2+\sigma_{np}^2)z+(\frac{\delta}{\phi})^{(\frac{1}{1+z})}(-(\sigma_p+\sigma_{np})^2+2(\sigma_p^2+\sigma_{np}^2)z)}{(\sigma_p+\sigma_{np})^3} & -\frac{\left((\frac{\delta}{\phi})^{\frac{1}{1+z}}\right)^{1+z}\sigma_{np}}{\sigma_p+\sigma_{np}} & -\frac{\left((\frac{\delta}{\phi})^{\frac{1}{1+z}}\right)^{1+z}\sigma_p}{\sigma_p+\sigma_{np}} \\
\frac{\phi\sigma_p\left(-2(\frac{\delta}{\phi}^{\frac{1}{1+z}}+\sigma_p+\sigma_{np})\right)(1+z)}{(\sigma_p+\sigma_{np})^2} & \frac{\phi\sigma_{np}\left(-2(\frac{\delta}{\phi}^{\frac{1}{1+z}}+\sigma_p+\sigma_{np})\right)(1+z)}{(\sigma_p+\sigma_{np})^2} & -\delta+\left(\frac{\delta}{\phi}^{\frac{1}{1+z}}\right)^{1+z}\phi & 0 \\
\frac{\phi\sigma_{np}\left(-2(\frac{\delta}{\phi}^{\frac{1}{1+z}}+\sigma_p+\sigma_{np})\right)(1+z)}{(\sigma_p+\sigma_{np})^2} & \frac{\phi\sigma_p\left(-2(\frac{\delta}{\phi}^{\frac{1}{1+z}}+\sigma_p+\sigma_{np})\right)(1+z)}{(\sigma_p+\sigma_{np})^2} & 0 & -\delta+\left(\frac{\delta}{\phi}^{\frac{1}{1+z}}\right)^{1+z}\phi
\end{pmatrix}$$

### Appendix A2: Linear functional response

When the enemy response to the victim is linear($z = 0$), we get the following set of eigenvalues.

$$\lambda_1 = \frac{(\sigma_p - \sigma_{np})\sqrt{\delta}\sqrt{2\delta - \phi(\sigma_{np} + \sigma_p)}}{\sqrt{\phi}(\sigma_{np} + \sigma_p)^{3/2}} \tag{7a}$$

$$\lambda_2 = -\frac{(\sigma_p - \sigma_{np})\sqrt{\delta}\sqrt{2\delta - \phi(\sigma_{np} + \sigma_p)}}{\sqrt{\phi}(\sigma_{np} + \sigma_p)^{3/2}} \tag{7b}$$

$$\lambda_3 = \frac{-(\sigma_p + \sigma_{np})\delta + \sqrt{\delta(\sigma_p + \sigma_{np})^2(\delta + 2\delta\phi(\sigma_p + \sigma_{np}) - \phi^2(\sigma_{np} + \sigma_p)^2)}}{\phi(\sigma_{np} + \sigma_p)^2} \tag{7c}$$

$$\lambda_4 = \frac{-(\sigma_p + \sigma_{np})\delta - \sqrt{\delta(\sigma_p + \sigma_{np})^2(\delta + 2\delta\phi(\sigma_p + \sigma_{np}) - \phi^2(\sigma_{np} + \sigma_p)^2)}}{\phi(\sigma_{np} + \sigma_p)^2} \tag{7d}$$

It is trivial to show that $\lambda_1$ and $\lambda_2$ are purely imaginary under the feasibility criteria of $\sigma_p + \sigma_{np} > 2\frac{\delta}{\phi}$. Let's investigate the sign of the real parts of $\lambda_3$ and $\lambda_4$, i.e. of $\Re(\lambda_3)$ and $\Re(\lambda_4)$. To simplify the analysis, let's make a small substitution $\sigma = \sigma_p + \sigma_{np}$. We get,

$$\lambda_3 = \frac{-\sigma\delta + \sqrt{\delta\sigma^2(\delta + 2\delta\phi\sigma - \phi^2\sigma^2)}}{\phi\sigma^2} = \frac{-\delta + \sqrt{\delta(\delta + 2\delta\phi\sigma - \phi^2\sigma^2)}}{\phi\sigma} \tag{8a}$$

$$\lambda_4 = \frac{-\sigma\delta - \sqrt{\delta\sigma^2(\delta + 2\delta\phi\sigma - \phi^2\sigma^2)}}{\phi\sigma^2} = \frac{-\delta + \sqrt{\delta(\delta + 2\delta\phi\sigma - \phi^2\sigma^2)}}{\phi\sigma} \tag{8b}$$

Since $\sigma > \frac{2\delta}{\phi}$ is required for feasibility, so is $\sigma\phi > 2\delta$ (since $\phi > 0$). This means there exists an $\epsilon > 0 \in \mathbb{R}$ such that $\epsilon = \sigma\phi - 2\delta$ for the feasibility criteria to be satisfied. From this, we can write

$$\lambda_{3,4} = \frac{-\delta \pm \sqrt{\delta(\delta + 2\delta\phi\sigma - \phi^2\sigma^2)}}{\phi\sigma} \tag{9a}$$

$$= \frac{-\delta \pm \sqrt{\delta(\delta + 2\delta(2\delta + \epsilon) - (2\delta + \epsilon)^2)}}{(2\delta + \epsilon)} \tag{9b}$$

$$= \frac{-\delta \pm \sqrt{\delta}\sqrt{\delta - 2\delta\epsilon - \epsilon^2)}}{2\delta + \epsilon} \tag{9c}$$

Our goal is to find if it is possible for $\Re(\lambda_i) > 0$ for $i = 3, 4$ in order to determine the stability of the system. Since $-\delta < 0$, this will only occur if the expression ($\sqrt{\delta}\sqrt{\delta - 2\delta\epsilon - \epsilon^2}$) is real and larger in magnitude than $-\delta$ and gets added to $-\delta$, i.e., $\lambda_3$. So, we need $-\delta + \sqrt{\delta}\sqrt{\delta - 2\delta\epsilon - \epsilon^2} > 0$. For this to happen, we need

$$\sqrt{\delta}\sqrt{\delta - 2\delta\epsilon - \epsilon^2} > \sqrt{\delta}\sqrt{\delta} \tag{10a}$$

$$\delta - 2\delta\epsilon - \epsilon^2 > \delta \tag{10b}$$

$$-2\delta\epsilon - \epsilon^2 > 0 \tag{10c}$$

But this will *not* occur for $\epsilon > 0$ (and $\delta > 0$) which we need for feasibility. Hence, for regions of feasible coexistence, $\Re(\lambda_{3,4}) \leq 0$.

As mentioned in the main text, the combination of the fact that $\lambda_{1,2}$) are pure imaginary and that $\Re(\lambda_{3,4}) \leq 0$ allow us to conclude that ReDEv$= 0$ in this case.

### Appendix A3: Eigenvalues across specificity and functional response

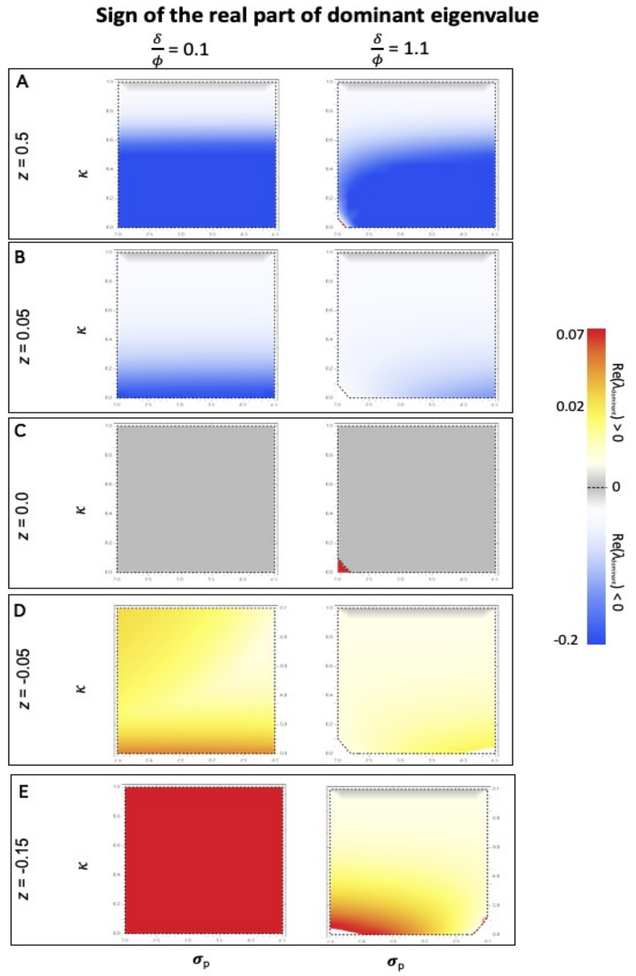


Figure A3.1: These heatplots shows the sign of the real part of the dominant eigenvalue (ReDEv) across functional responses and enemy specificity. The susceptibility of the preferred victim to its enemy varies on the x-axis. On the y-axis the ratio of non-preferred susceptibility to preferred susceptibility ($\kappa = \frac{\sigma_{np}}{\sigma_p}$) varies from 0 to 1, i.e. from specialist enemies at the bottom of the plot to generalist enemies at the top. Black dashed line encloses the feasible coexistence region. Note that ReDEv is $< 0$ almost everywhere for accelerating functional responses ($z > 0$) (the exception being complete generalist enemies), ReDEv $= 0$ so long as there is a feasible equilibrium with a linear functional response ($z = 0$), and ReDEv $> 0$ almost everywhere for saturating functional responses ($z < 0$) (the exception being complete generalist enemies).

### Appendix A4: Accelerating functional response numerical simulations

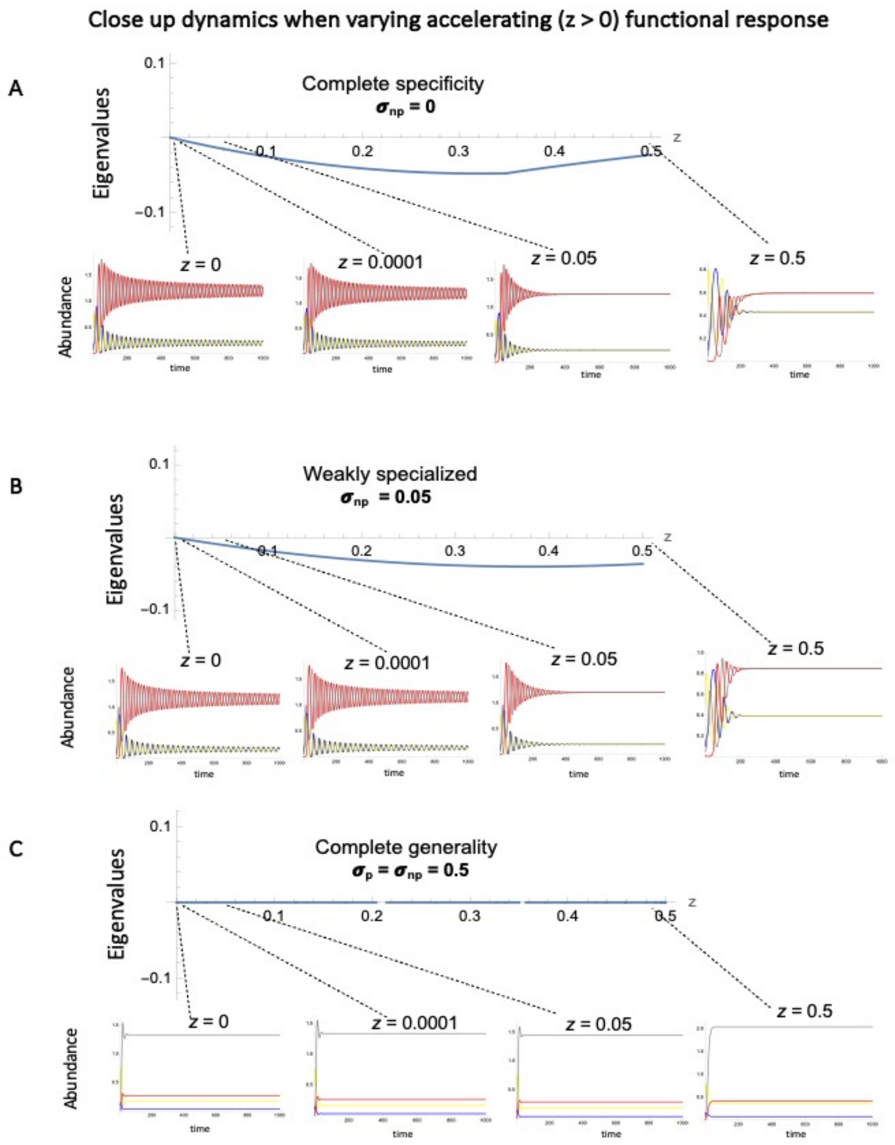


Figure A4.1: A closer look at the dynamics as we vary the degree of acceleration in the accelerating functional response from Fig.4(A)(main text). We can see that for both specificity in (A)(Fig.3 slice 6, main text) and weak generalization in (B)(Fig.3 slice 8, main text), the system approaches the equilibrium point faster the larger $z$ is. For complete generalists in (C)(Fig.3 slice 7, main text), it appears to reach a fixed point very quickly; but in fact, what we are capturing is a very long period length where one enemy-victim pair appears to settle down at reasonable density where as the other pair persists near zero densities. Other variables: $\delta = 0.1, \phi = 1, \sigma_p = 0.5$

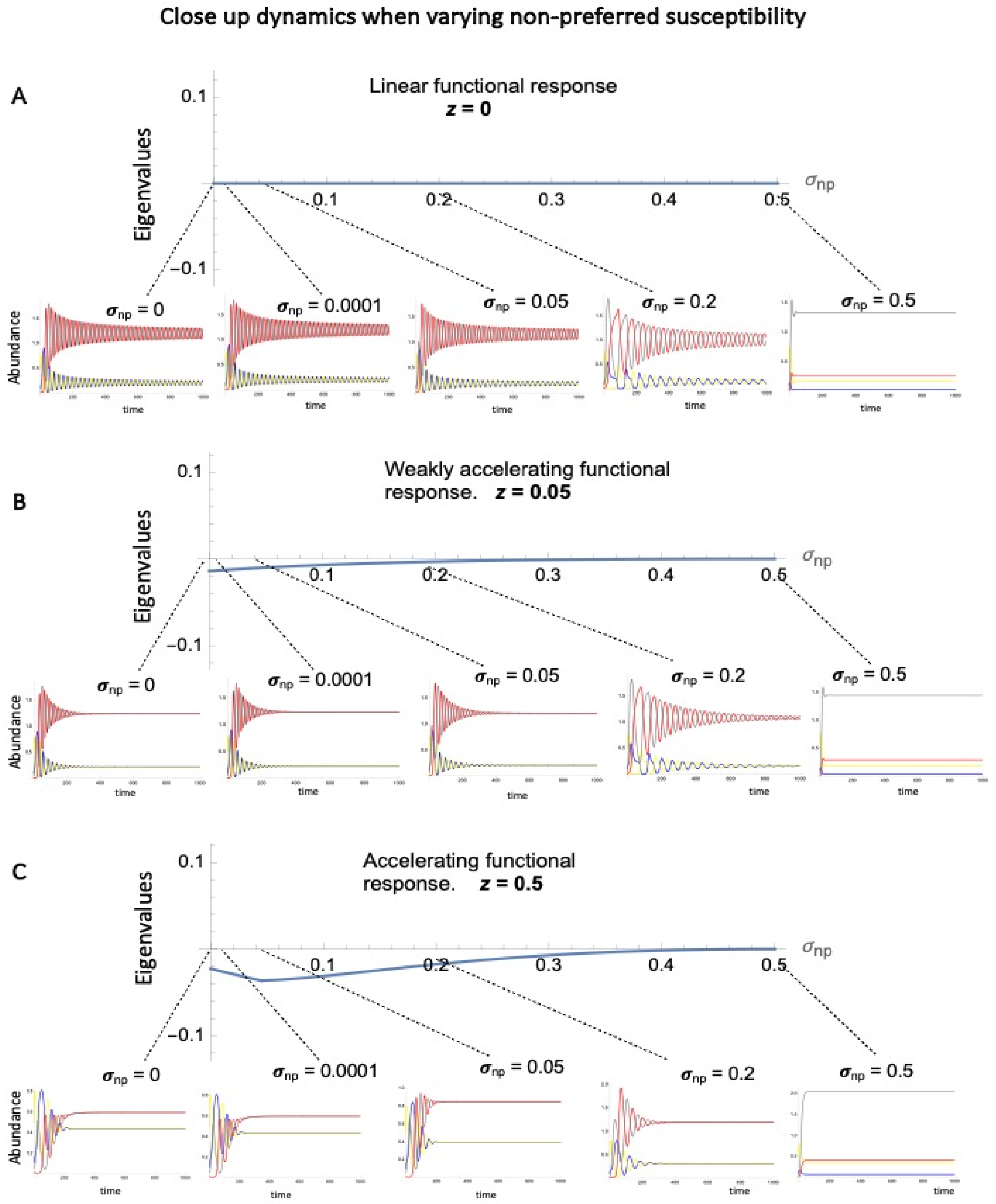


Figure A4.2: A closer look at the dynamics as we vary the host specificity in the case of accelerating functional responses from Fig4(B)(main text). We see that for all values of $z$ the system approaches the equilibrium point more slowly the weaker the specificity (i.e. higher $\sigma_{np}$). For each of these cases where $\sigma_n p = \sigma_p = 0.5$, we see the same behaviour as the previous Fig. A5.1. (A) corresponds to slice 11, Fig.3, main text. (B) corresponds to slice 9, Fig.3, main text. (C) corresponds to slice 10, Fig.3, main text. Other variables: $\delta = 0.1, \phi = 1, \sigma_p = 0.5$

### Appendix A5: Saturating functional response

Here we provide additional details on the properties and stability of dynamical features shown in the main text.

*Effects of varying functional response non-linearity*

For $z = -0.07$, we see limit cycles of Type 2; these are symmetric limit cycles with one host-pathogen pair exhibiting dynamics that are identical to the other, but shifted by half a period. As $z$ decreases, these Type 2 limit cycles eventually lose stability, giving rise to limit cycles of Type 1. Individual Type 1 limit cycles are not symmetric: one host-pathogen pair exhibits temporal dynamics that are different than the other, and not just by a phase shift. This may initially be surprising, given that the underlying ODE model is perfectly symmetric. The explanation lies in the fact that Type 1 limit cycles actually exist in pairs, where the roles of PA-H1 and PB-H2 are reversed (see example for $z = -0.16$). These represent alternative, stable limit cycles, separated in 4D space by the existence of the unstable Type 2 limit cycle that exists in this region. This is partially visible in the phase plane diagram for this $z$ value (compare blue dashed lines with solid orange lines). As $z$ continues to decrease, the Type 1 limit cycles also lose stability, giving rise to a region of parameter space where there are 3 simultaneous unstable limit cycles; this produces chaotic dynamics (see $z = -0.17$). For a small range of more extreme $z$ values, the Type 2 limit cycle briefly regains stability (see $z = -0.1815$), before disappearing. Beyond this range, only the unstable Type 1 limit cycles still exist, and the overall system loses stability, exhibiting increasingly extreme oscillations that eventually drive one host or the other to extinction, after which the coexistence of a single host-pathogen pair remains possible for a while (see main text).

*Effects of varying specificity*

As with variation in $z$, changing $\sigma_{np}$ values gives rise to a series of distinct limit cycles and behaviors which we expand upon here (Fig. A5.2). Starting at $\sigma_{np} = 0.05$ (for $z = -0.15$; this corresponds to the intersection of slices 3 and 5 in Fig. 3 of the main text), we again note that two types of limit cycles exist with this parameterization: a pair of asymmetric stable limit cycles (Type 1 in Fig. A5.2) and an unstable, symmetric limit cycle (Type 2 in Fig. A5.2). As $\sigma_{np}$ increases from there (corresponding to less specificity), the Type 1 limit cycle collapses into the Type 2 limit cycle, which becomes stable (see $\sigma_{np} = 0.09$, Fig. S2). We then see the emergence of a new set of limit cycles with a longer period that represent an alternate stable outcome for the system (e.g., at $\sigma_{np} = 0.12$, Type 2 and Type 3 both exist, Fig. A5.2). These are connected to each other by a branch of unstable limit cycles (black dashed line). This pattern repeats several more times as $\sigma_{np}$ continues to increase, giving rise to limit cycle Types 4, 5, and eventually 6, as period length continues to increase (Fig. I, bottom row). These are also

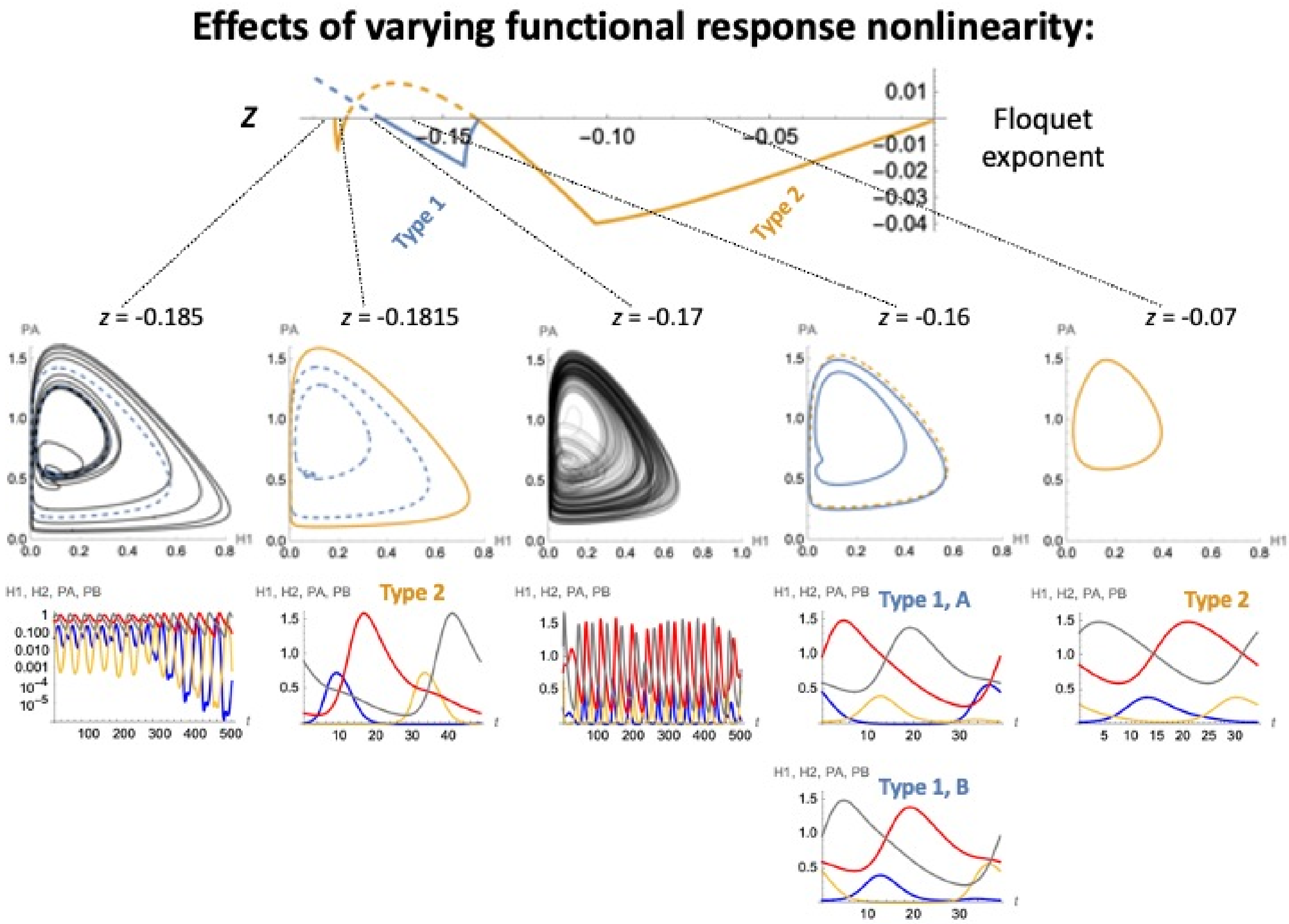


Figure A5.1: A closer look at the dynamics as we vary the saturating functional response. Details on dynamical features emerging as $z$ varies, for $\sigma_{np} = 0.05$ (slice 3 in Fig.3, and bottom row of Fig.5). Two distinct types of limit cycles exist over this range (Type 1 in blue, Type 2 in orange). The top panel illustrates the stability of these limit cycles across varying z values, with stable (or unstable) limit cycles indicated by solid (or dashed) lines and dominant Floquet exponents $< 0$ (or $> 0$). Complementing this overview, specific examples of these limit cycles, chaos, and extinction are shown in detail in the lower panels at particular values of $z$, including phase planes (for a focal host H1 and pathogen PA) in the top row, and temporal dynamics of all four species in the bottom row ($H1$ = blue, $H2$ = green,$PA$ = red, $PB$ = pink).

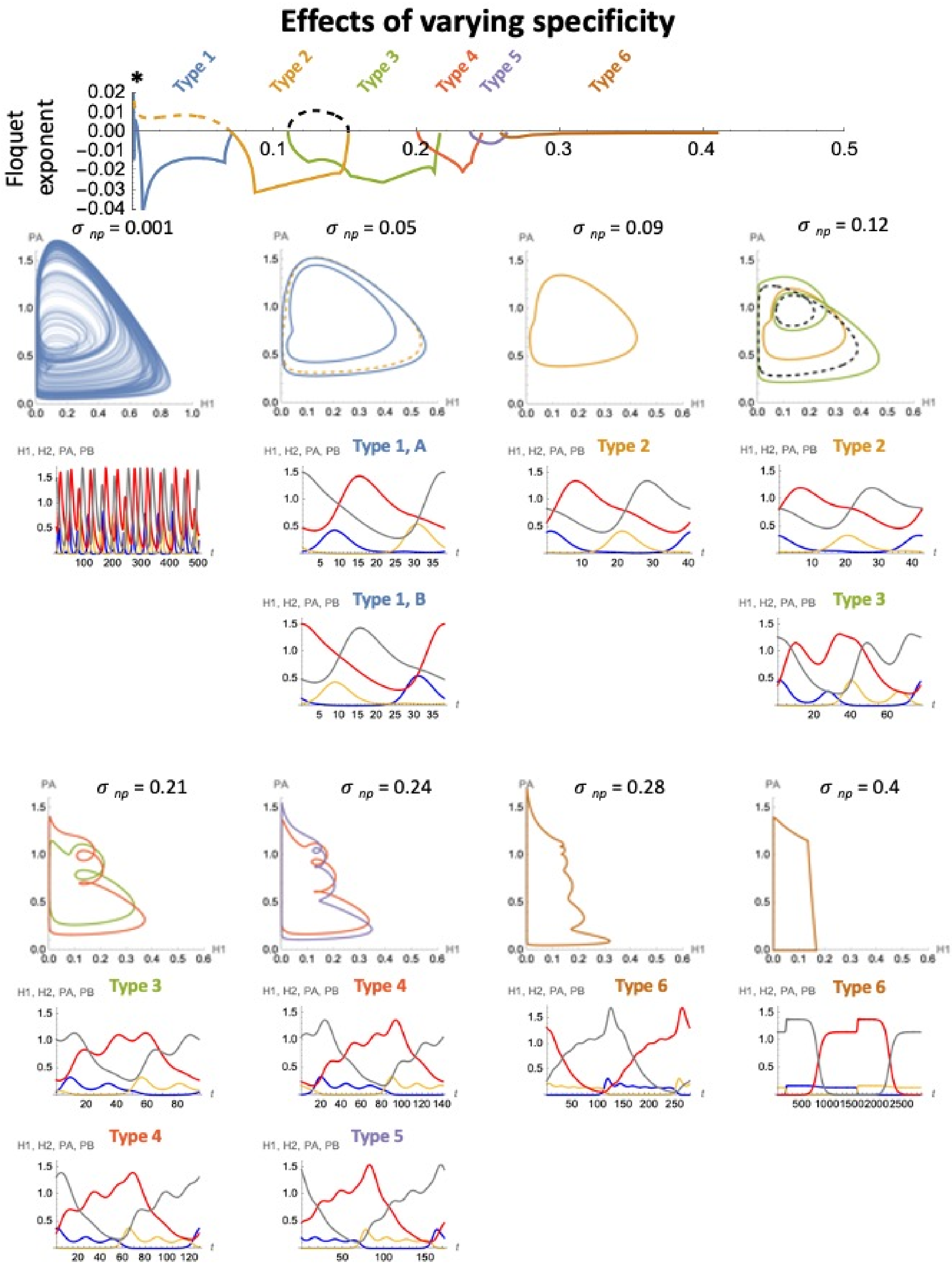


Figure A5.2: Details on dynamical features emerging as $\sigma_{np}$ varies, for $z = -0.15$ (slice 5 in Fig.3, main text, and bottom row of Fig.5 I). At least seven distinct types of limit cycles exist over this range, labeled Types 1-7 and distinguished by separate colors in the top panel, which shows variation in the dominant Floquet exponent associated with each type. As in Fig. A5.1, we indicate stable (or unstable) limit cycles by solid (or dashed) lines, corresponding to dominant Floquet exponents $< 0$ (or $> 0$). Complementing this overview, specific examples of these limit cycles and intervals of chaos are shown in detail in the lower panels at particular values of $\sigma_{np}$, including phase planes (for a focal host $H1$ and pathogen $PA$) in the top row, and temporal dynamics of all four species in the bottom row ($H1$ = blue,$H2$ = green, $PA$ = red, $PB$ = pink). Alternative stable limit cycles exist for multiple regions of this parameter space; in these cases, their dynamics are shown within the same phase plane to facilitate comparison, or labeled/colored accordingly. Several additional features emerge for very small values of $\sigma_{np}$ (marked by the *); these are highlighted in a subsequent figure.

likely to be connected to each other via sets of unstable limit cycles, but these are difficult to find numerically. After the onset of the Type 6 cycles, further increases in $\sigma_{np}$ do not yield new types of limit cycles; however, we do observe dramatic increases in the period of fluctuation of limit cycle solutions, which eventually approach infinite length as $\sigma_{np}$ approaches $\sigma_p = 0.5$. This is characteristic of an infinite period bifurcation ( Strogatz 2018). Next we turn to considering how the system changes as $\sigma_{np}$ decreases from 0.05, taking us in the other direction from the intersection of slices 3 and 5 (Fig. 3). A number of features occur within a compressed region of parameter space, which we expand on in Figure A5.3 below. In particular, we see the paired asymmetric limit cycles (Type 1) briefly lose stability and are replaced by a different pair of asymmetric limit cycles with a longer period (see $\sigma_{np} = 0.003$ in Fig. A5.3), which we label Type 7. These persist for only a small range of $\sigma_{np}$ values, before they collapse again back into stable limit cycles of Type 1, which then proceed to quickly lose stability, yielding chaotic oscillations (e.g., $\sigma_{np} = 0.001$). Chaotic solutions extend until $\sigma_{np}$ values are very nearly zero ($< 10^{-8}$), at which point the four species system collapses as one host is driven to extinction (see main text). Figure A5.3. Details on variation in the model's dynamical features over a range of small $\sigma_{np}$ values, for $z = -0.15$ (slice 5 in Fig. 3, and bottom row of Fig. 5, main text). Layout, details, and other parameters are as described in Fig. A5.2. This figure shows in particular the emergence of the Type 7 limit cycle at $\sigma_{np} = 0.003$. For extremely small values of $\sigma_{np}$ ($< 10^{-8}$), the four species system collapses (not shown).

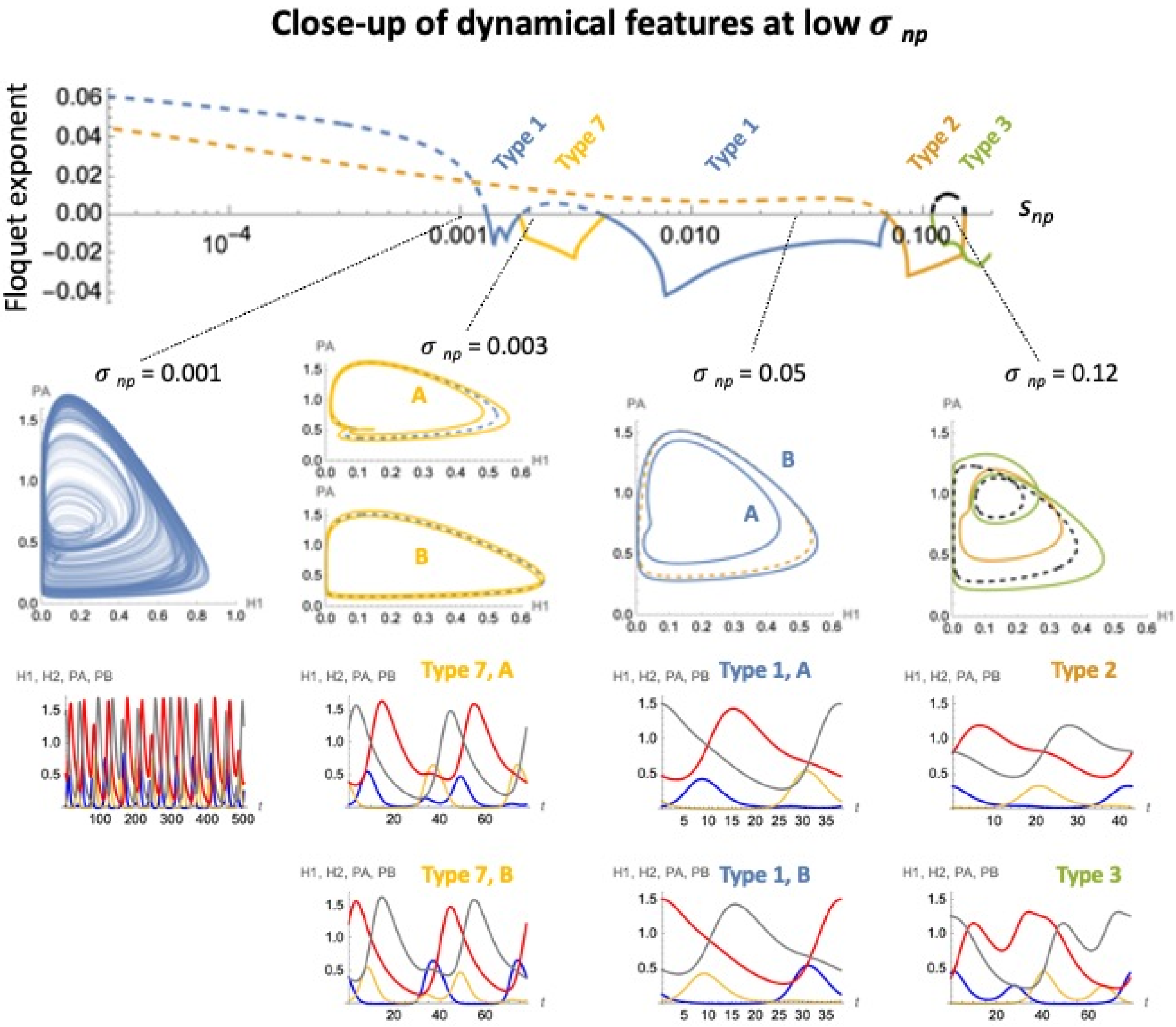


Figure A5.3: Details on variation in the model's dynamical features over a range of small $\sigma_{np}$ values, for $z = -0.15$ (slice 5 in Fig.3, and bottom row of Fig.5). Layout, details, and other parameters are as described in Fig. A5.2. This figure shows in particular the emergence of the Type 7 limit cycle at $\sigma_{np} = 0.003$. For extremely small values of $\sigma_{np}$ (less than $10^{-8}$), the four species system collapses (not shown).
